# Failed apoptosis enhances melanoma cancer cells aggressiveness

**DOI:** 10.1101/755744

**Authors:** K. Berthenet, C. Castillo Ferrer, N. Popgeorgiev, H. Hernandez-Vargas, G. Ichim

## Abstract

Triggering apoptosis remains an efficient strategy to treat cancer. However, apoptosis is no longer a final destination, since cells can undergo partial apoptosis without dying. Recent evidence shows that partial mitochondrial permeabilization and non-lethal caspase activation occur under certain circumstances, though it remains unclear how failed apoptosis impacts established cancers. Using a cancer cell model to trigger non-lethal caspase activation based on either BH3-only protein expression or chemotherapy treatment, we found that melanoma cancer cells failing to undergo complete apoptosis have a particular transcriptomic signature associated with focal adhesions, transendothelial migration and modifications of actin cytoskeleton. In line with this, cancer cells surviving apoptosis have a gain in migratory and invasive properties both *in vitro* (random migration, chemotaxis, wound healing and invasion assays) and *in vivo* (in a model of zebrafish metastasis) We further demonstrate that the failed apoptosis-associated gain in invasiveness is regulated by the c-Jun N-terminal kinase (JNK) pathway while its RNA seq signature is found in metastatic melanoma. These findings are highly significant for understanding how cell death can both cure and promote cancer.

## Introduction

Cell death and more specifically apoptosis is without a doubt the spearhead of many anti-cancer therapies, ranging from conventional chemotherapy and radiotherapy to recently developed targeted therapy and immunotherapy ^1^ ^2^. In general, apoptosis is triggered either by the engagement of death receptors, including TNFR or FAS (the extrinsic pathway), or by an intracellular death signal, such as DNA damage, tumor suppressor activation, nutrient deprivation or ER stress (the intrinsic pathway). It is commonly accepted that the point of no return in the apoptotic process is the mitochondrial outer-membrane permeabilization (MOMP), followed by rapid loss of mitochondrial function, cytochrome *c* release, apoptosome assembly and activation of caspase-9 prior to effector caspase-3 and −7 ^3^ ^4^. Given its central role in the apoptotic process, MOMP is tightly regulated by BCL-2 family proteins that are either anti-apoptotic (BCL-2, BCL-xL or MCL1), pro-apoptotic (BID, BIM, BAD, PUMA or NOXA), or effector pore-forming proteins (BAX and BAK) ^5^.

MOMP ensures efficient effector caspase activation and kills a cell within minutes by cleaving hundreds of vital protein substrates ^4^. However, recent research has consistently shown that exceptions to this rule are not uncommon, as MOMP can either be incomplete, with a few intact mitochondria overcoming wide-spread permeabilization and serving as “seeds” for mitochondrial re-population (the incomplete MOMP scenario), or occur in only a fraction of mitochondria (minority MOMP) when lethal stress is of low amplitude ^6^ ^2^ ^7^ ^8^. On one hand, these scenarios explain how caspases may be activated at non-lethal levels and have vital roles in various physiological processes such as macrophages differentiation, muscle and neuronal function and even stemness or induced pluripotent stem cells ^9^ ^10^ ^11^ ^12 13 14^. On the other hand, failed apoptosis could have a dark side by damaging the DNA, triggering genomic instability and favoring oncogenic transformation ^15^ ^8^. Despite the importance of cell death in physiological settings and disease, there is still very little scientific evidence of the phenotypic consequences of failed apoptosis on cancer cells. This is particularly important since drugs used in cancer management do not kill all cancer cells and those surviving are likely responsible for cancer relapse, drug resistance, metastasis and increased mortality.

This study therefore aimed at assessing the phenotypic effects of failed apoptosis. For this, we focused on melanoma since it is one of the most aggressive cancers, with a high rate of mortality. Using a sensitive caspase activation reporter in settings of failed apoptosis, we isolated and thoroughly characterized melanoma cancer cells surviving the partial induction of apoptosis. Importantly, our results suggest that these cells gain in aggressiveness, displaying increased migratory and invasive potentials, governed by the JNK-AP1 transcriptional axis.

## Results

### Establishment of a melanoma cellular model for triggering and isolating cells undergoing failed apoptosis

To assess the effects of failed apoptosis on the phenotype of surviving cells, we used a BiFC-based (bi-molecular fluorescence complementation) caspase reporter assay that provides a direct approach for the efficient visualization and quantification of minute amounts of active effector caspases ^16^ ^7^ (Figure 1A). This reporter was therefore expressed *via* lentiviral transduction into various human melanoma cell lines with different metastatic potentials, and its efficacy was first validated by flow cytometry (Figure 1B, C and S1A, B for additional cell lines). We then added a doxycycline-inducible system controlling the expression of the pro-apoptotic truncated BID (tBID) protein, capable of activating BAX and thus inducing rapid MOMP and cell death (Figure 1A). As shown in Figure 1D and S1C, D for 3 melanoma cell lines, doxycycline treatment at a high dose (1 μg/mL) induced tBID expression and cleavage of PARP1, indicative of the induction of apoptosis. Importantly, the pan-caspase inhibitor Q-VD-OPh blocked caspase activation. We then sought to incrementally reduce the concentration of doxycycline in order to obtain failed apoptosis, characterized by partial MOMP and non-lethal caspase activation (see schematic diagram in Figure 1A). More specifically, WM115 cells were treated with doxycycline, ranging in concentration from 1 μg/mL to 10 ng/mL and the expression of tBID was validated by both western blot and immunofluorescence analyses (Figure 1E and S1E). Data suggest that tBID induction is proportional to doxycycline concentration. Regarding induction of cell death, while 1 μg/mL of doxycycline induced complete apoptosis, lower concentrations of 100 or 50 ng/mL killed only a fraction of melanoma cells, as determined by propidium iodide staining and IncuCyte-based live-cell microscopy in WM115 and WM852 cells (Figure 1F, G and S1F, G).

**Figure 1.**
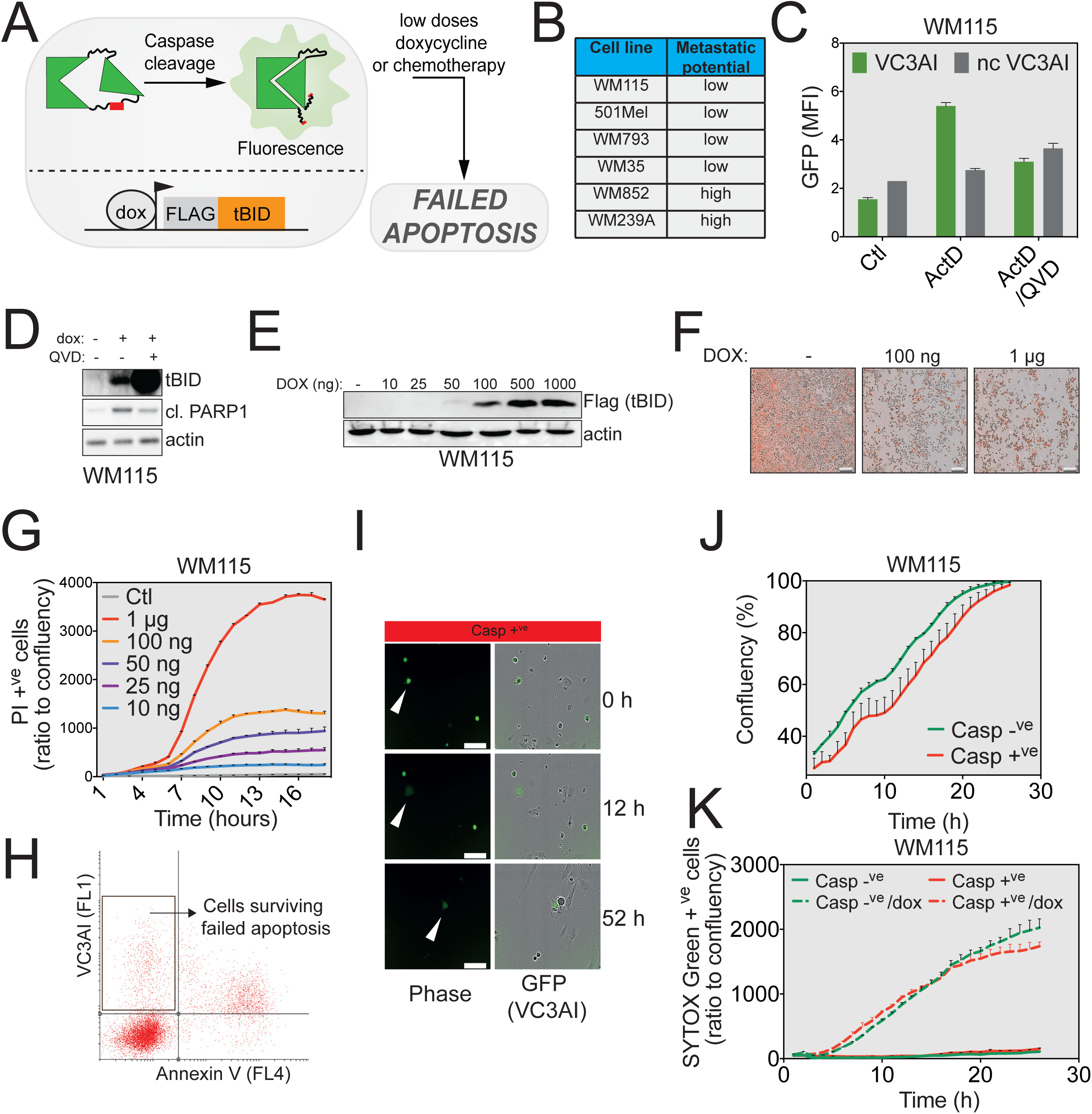
Generation and validation of the cellular model of failed apoptosis. **A**. Melanoma cancer cells generated to stably express the VC3AI caspase reporter together with a doxycycline-inducible tBID expression vector. Following low-dose doxycycline or chemotherapy treatment, this model mimics failed apoptosis. **B**. List of melanoma cell lines used in the present study and their metastatic potential. **C**. WM115 cells were treated with 1 µM actinomycin D (ActD) in the presence or absence of 10 µM Q-VD-OPH (QVD) for 24 hours. Caspase activation was then determined by flow cytometry. WM115 stably transfected with a non-cleavable form of VC3AI (ncVC3AI) served as control. **D**. Immunoblot analysis of tBID expression and cleaved PARP1 in WM115 cell line treated with 1 µg/mL of doxycycline with or without QVD for 24 hours. Actin served as a loading control. **E**. Titration of doxycycline concentration in WM115 cells. tBID expression was determined by western blot while actin served as a loading control. **F**. Representative images of propidium iodide (PI) staining of WM115 cells after 24 hours doxycycline treatment (100 ng or 1 µg) acquired using the IncuCyte ZOOM imaging system. **G**. WM115 cells were treated with various doses of doxycycline and apoptosis induction was measured in real time using PI staining and the IncuCyte ZOOM imaging system. **H**. Gating strategy used to sort and further characterize cells undergoing failed apoptosis (VC3AI^+ve^/AnnexinV^−ve^). **I**. Representative images of sorted VC3AI^+ve^ /AnnexinV^−ve^ WM852 cells immediately after, 12 hour and 52 hours post-sorting. **J**. Cell proliferation was determined using the IncuCyte ZOOM live-cell imaging based on the confluency parameter. **K**. Responsiveness to tBID-expression challenge was measured using SYTOX Green staining.

We then used this model to isolate by flow cytometry melanoma cells undergoing failed apoptosis. For this, WM115 or WM852 cells were treated with low doses of doxycycline, and 24 hours post-treatment VC3AI^+ve^/DAPI^−ve^/AnnexinV^−ve^ cells were sorted (Figure 1H for the gating strategy). This population, generically called caspase-positive cells (or simply Casp^+ve^ henceforth) is enriched in cells that trigger caspase activation at a non-lethal level, which is compatible with cell survival (hereafter called failed apoptosis). Of note, for each batch of Casp^+ve^ cells, we also sorted the VC3AI^−ve^/DAPI^−ve^/AnnexinV^−ve^ cells (Casp^−ve^), which is the counterpart population unaffected by cell death stimuli. These two populations were further characterized throughout this study. Although the Casp^+ve^ cells took more time to recover, three days post-sorting the caspase reporter VC3AI was still visible by immunofluorescence, while effector caspase activation was validated using a fluorometric assay based on detecting the cleavage of DEVD-AFC (7-amino-4-trifluoromethyl coumarin) (Figure 1I and S1H). When fully recovered, Casp^+ve^ and Casp^−ve^ cells had comparable proliferation rates and were equally sensitive to tBID-induced apoptosis (Figure 1J, K and S1I, J). This suggests that our cell sorting strategy did not favor the isolation or enrichment in a cell population generally resistant to apoptosis. We have thus set up an efficient melanoma cellular model to specifically trigger and isolate cells undergoing failed apoptosis.

### Failed apoptosis triggers a specific transcriptional program centered on cell motility

To define the impact of failed apoptosis at the molecular level and gain insight into specific cellular processes that might be affected, we performed RNA sequencing on both Casp^+ve^ and Casp^−ve^ populations originating from two different melanoma cell lines (WM115 and WM852 cells). Principal component analysis (PCA) showed that for both cell lines the Casp^−ve^ and Casp^+ve^ cells clearly form different clusters (Figure 2A). When plotting the fold change in gene expression according to their up- or down-regulation, we noticed that WM852 melanoma cells undergoing failed apoptosis (Casp^+ve^ cells) present more up-regulated than down-regulated genes, advocating for a specific transcriptional program associated with failed apoptosis (Figure 2B). Gene expressions obtained are further detailed in Figure 2C and S2A and B for the WM115 cell line, while validation of our RNASeq data was conducted by qRT-PCR on 5% randomly chosen genes within the total number of genes differentially expressed (Figure S2C).

**Figure 2.**
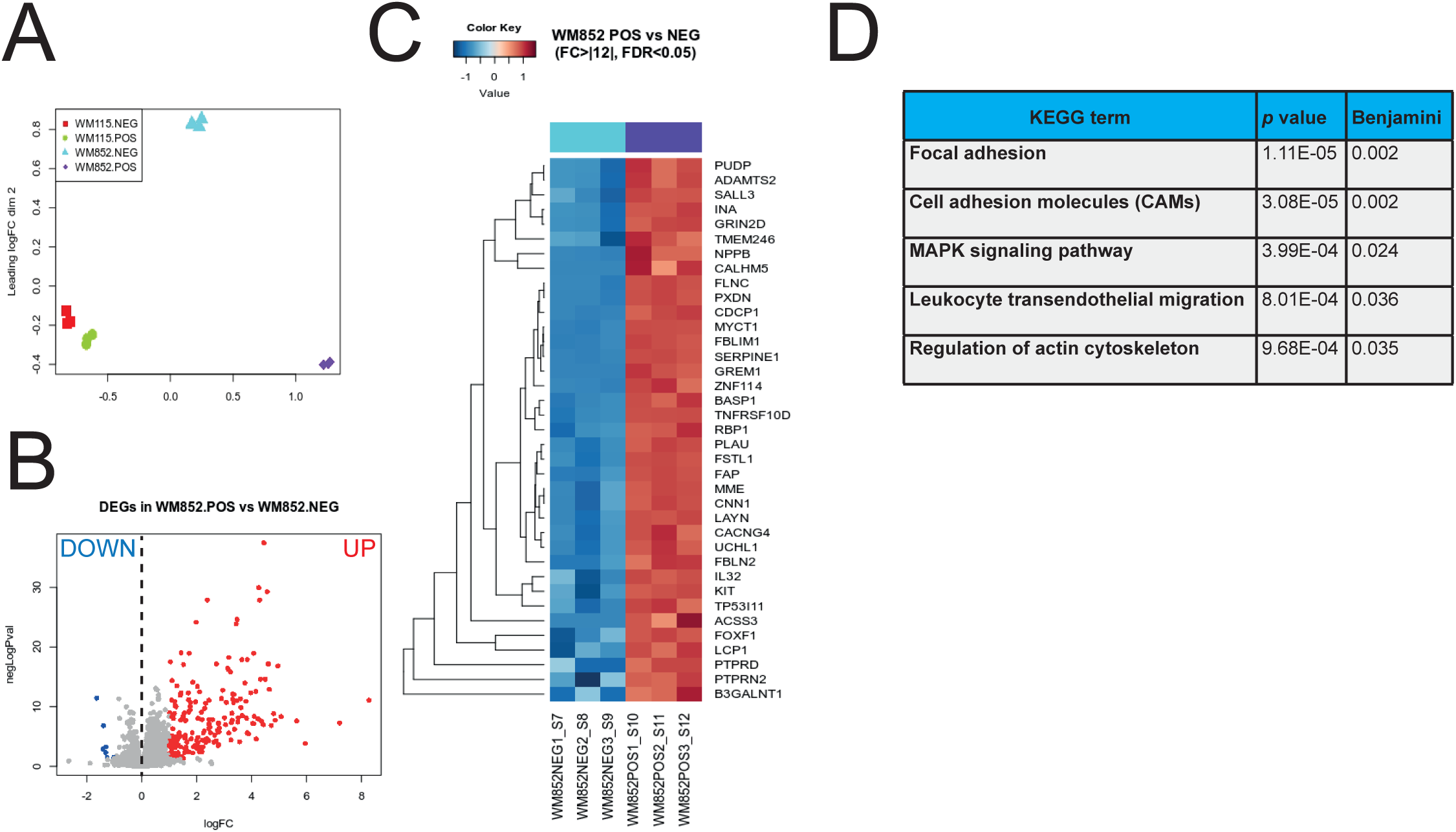
RNAseq highlights a transcriptional signature specific for failed apoptosis. **A**. PCA analysis of RNAseq data reveals clustering of Casp^−ve^ and Casp^+ve^ cells for both WM115 and WM852 cells. The analysis was conducted in triplicate. **B**. Vulcanoplot displaying the expression (in log fold change) of each gene differentially expressed in WM852 Casp^+ve^ compared to WM852 Casp^−ve^ cells. **C**. Unsupervised clustering of the RNAseq data in WM852 Casp^−ve^ and Casp^+ve^ cells. Red indicates increased while blue indicates decreased mRNA abundance of selected genes with fold change above 2. **D**. KEGG pathway enrichment analysis of the genes up-regulated following failed apoptosis in WM852 cells.

To identify pathways overrepresented in Casp^+ve^ cells compared to control cells, we performed a pathway analysis using the Kyoto Encyclopedia of Genes and Genomes (KEGG) *via* the DAVID database (https://david.ncifcrf.goV). For this we used WM852 cells that displayed the highest differences in PCA (Figure 2A). The most striking result that emerged from this analysis is that Casp^+ve^ cells present a transcriptional signature associated with focal adhesion, cell adhesion molecules, transendothelial migration or regulation of actin cytoskeleton (Figure 2D and S2D-G). Even though this was not as obvious in WM115 cells, WM852 and WM115 Casp^+ve^ cells have in common several up-regulated genes, all involved in cell motility (Figure S2H). These results suggest that failed apoptosis may have an impact on the melanoma cells capacity to migrate and invade.

### Cancer cells surviving apoptosis have an increased focal adhesion size and random migratory capacity

Given the gene signature centered on cell motility, we next evaluated the capacity of melanoma cells surviving failed apoptosis to adhere to extracellular matrix. To this end, WM852 cells (Casp^−ve^ and Casp^+ve^ populations) were seeded onto matrigel-coated culture dishes and imaged for 24 hours using an IncuCyte live-cell imager. As shown in representative pictures taken 1 hour and 24 hours after seeding, melanoma cells undergoing failed apoptosis exhibited a better adhesion to the matrix, compared to control cells (Figure 3A). This was quantified over time and these findings were substantiated in WM852 (Figure 3B) and WM793 cells (Figure S3A, B). Cells adhere to their respective substrates through protein complexes called focal adhesions that link the intracellular cytoskeleton to the extracellular matrix via integrins ^17^. Aside from their role as anchors, focal adhesions are also involved in cell migration, especially during metastasis ^18^. To correlate enhanced matrix adhesion with focal adhesion size in our cells, we used immunofluorescence to highlight two key components of focal adhesions, namely paxillin and vinculin. In cells surviving apoptosis, the size of focal adhesions was significantly larger than that of Casp^−ve^ cells, as evidenced both in WM852 (Figure 3C-F) and WM793 cells (Figure S3C-F). Since focal adhesions can also mediate cell migration depending on the rate of their turnover, we next wondered whether Casp^+ve^ cells have enhanced migratory properties. To test this possibility, we first performed a random migration assay in which cancer cells were seeded at a very low density and we then used IncuCyte-based live-cell microscopy to quantify their random non-directional migration over time. As shown in the representative spider plots (Figure 3G) and associated histogram (Figure 3H), melanoma cells surviving the initiation of apoptosis had a greater non-directional migratory capacity. Since parental cells behaved as Casp^−ve^ cells in most of our experiments, we henceforth compared only Casp^−ve^ and Casp^+ve^ cells. Taken together, these results suggest that there is an association between the transcriptional signature and the migratory capacity of melanoma cancer cells failing to achieve apoptosis.

**Figure 3.**
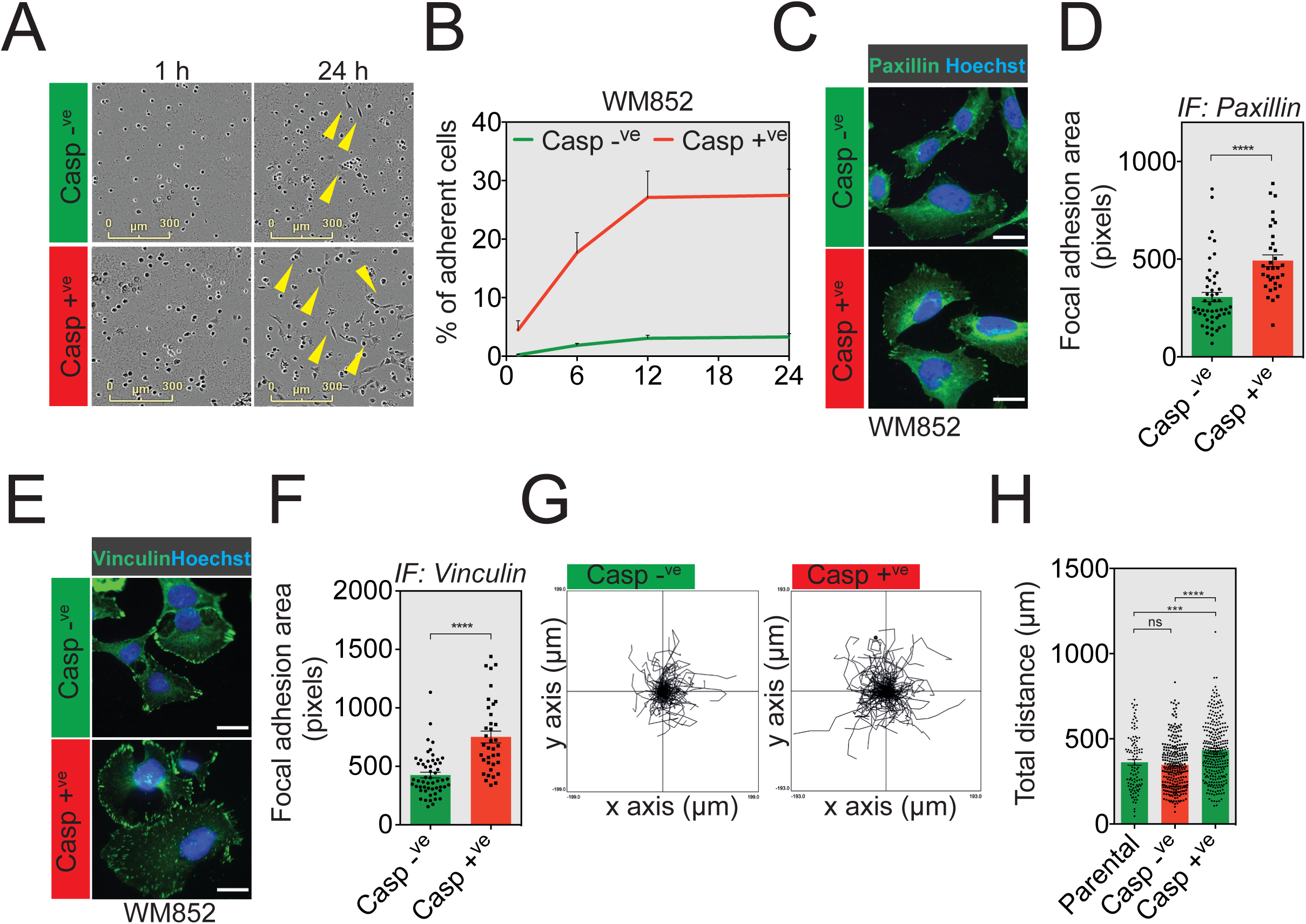
Failed apoptosis promotes cell adhesion. **A**. Casp-^ve^ and Casp^+ve^ WM852 melanoma cells were seeded onto a 96-well plate previously coated with 100 µg/mL matrigel and imaged using the IncuCyte ZOOM system. Representative images of the adhesion capacity of WM852 are shown at 1 hour and 24 hours after seeding. **B**. Quantification of the number of WM852 adherent cells. **C**. Paxillin immunostaining analysis in Casp^−ve^ and Casp^+ve^sorted WM852 melanoma cells. **D**. Quantification of focal adhesion area based on paxillin immunostaining. **E**. Vinculin immunostaining analysis in Casp^−ve^ and Casp^+ve^ sorted WM852 melanoma cells. **F**. Quantification of focal adhesion area based on vinculin immunostaining. **G**. Casp^−ve^ and Casp^+ve^ WM852 melanoma cells were seeded onto an ImageLock 96-well plate at low density and imaged for 24 hours. Each spider graph is a composite of the migratory paths of several individual cells. **H**. Quantification of total distance of random migration between parental, Casp^−ve^ and Casp^+ve^ sorted WM852 melanoma cells.

### Failed apoptosis contributes to the migratory and invasive potential of melanoma cells

Chemotaxis and directional migration and invasion through the extracellular matrix are essential for metastatic dissemination ^18^. The next set of experiments focused on determining whether failed apoptosis enhances tumor cell directional migration and invasion. We initially tested the migration of Casp^−ve^ and Casp^+ve^ WM115 cells seeded in serum-free medium in the upper compartment of a Boyden chamber towards the lower chamber filled with 20% serum-containing medium (Figure 4A for experimental setup), by staining the migrating cells with vital Hoechst (Figure 4B for representative images). As shown in Figure 4C, Casp^+ve^ WM115 cells display a chemotactic advantage compared to Casp^−ve^ cells. This result was validated using WM115 cells surviving treatment with the BH3-mimetic ABT-737, capable of inducing failed apoptosis, and further corroborated using WM793 cells (Figure S4A-D). Next, we conducted an Incucyte live-imager based wound-healing assay to monitor and quantify directional cancer cell migration over time (Figure 4D).

**Figure 4.**
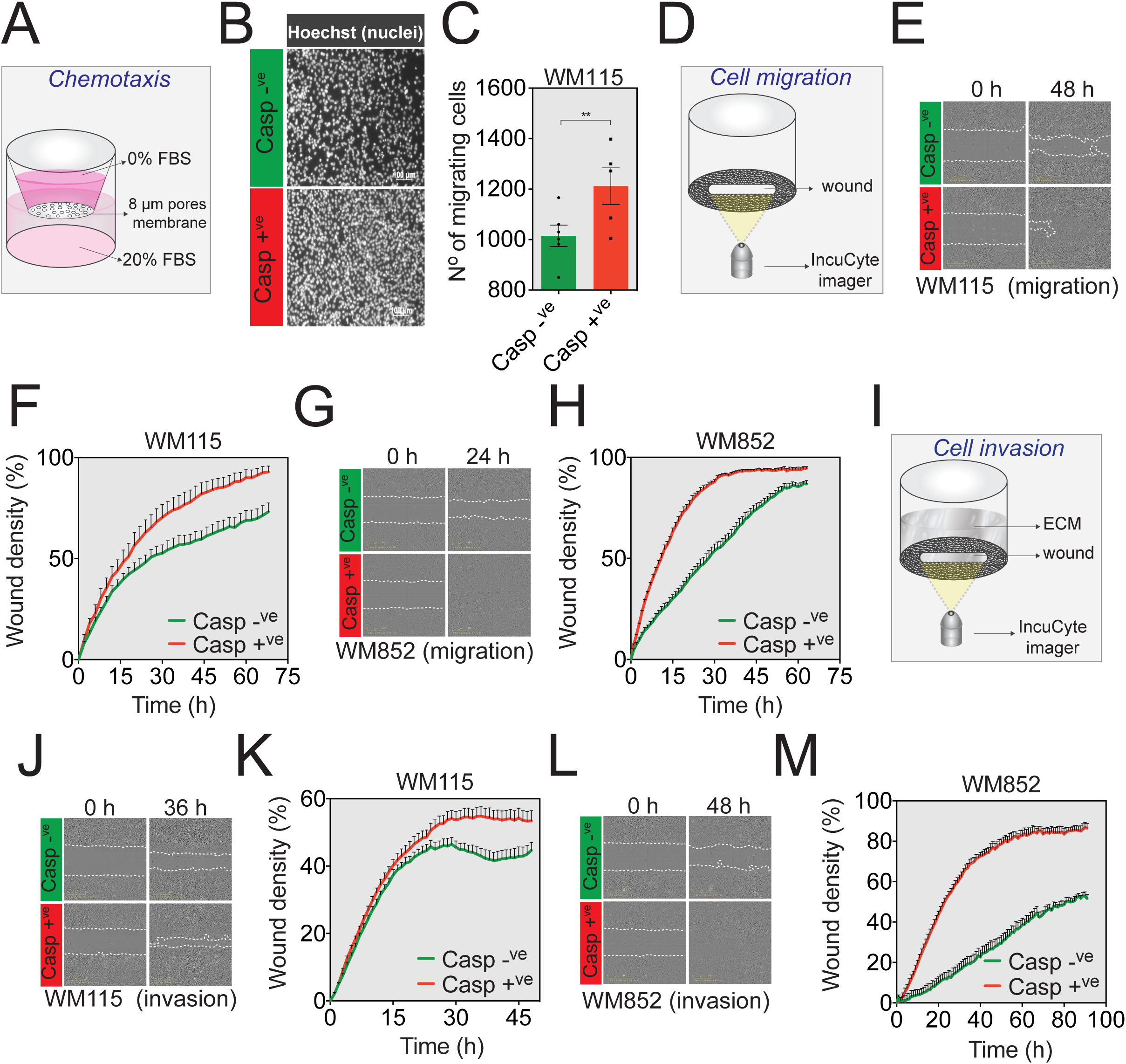
Melanoma cell lines surviving apoptosis acquire an aggressive potential. **A**. Schematic representation of the chemotaxis assay based on the capacity of melanoma cells to migrate through 8-µm pores from serum-depleted medium towards 20% serum-containing culture medium. **B**. Representative images of sorted Casp^−ve^ and Casp^+ve^ WM115 cells stained with Hoechst that crossed the transwell membrane. **C**. Quantification of sorted WM115 having crossed the transwell membrane. **D**. Schematic representation of the IncuCyte ZOOM-based wound-healing assay relying on the capacity of melanoma cells to migrate and close the wound. **E**. Monolayer WM115 cells were wounded and photographs were taken immediately after wound induction and 48 hours later. **F**. IncuCyte ZOOM-based quantification of the migratory potential of sorted WM115 cells through wound area measurement. **G**. Monolayer WM852 cells were wounded and photographs were taken immediately after wound induction and 24 hours later. **H**. Quantification of sorted WM852 migration potential through wound area measurement using the IncuCyte ZOOM live-cell imager. **I**. Schematic representation of the IncuCyte ZOOM-based invasion assay relying on the capacity of melanoma cells to invade through a layer of matrigel and close the wound. **J**. Representative images of the wound in a layer of sorted WM115 at 0 and 36 hours after wounding in the presence of matrigel. **K**. IncuCyte ZOOM-based measurement of WM115 cells invasion through matrigel. **L**-**M**, The same analyses as in **J** and **K** were performed using WM852 cells.

As depicted in the representative images (Figure 4E) and clearly evidenced in the time-course wound healing quantification plot (Figure 4F), Casp^+ve^ WM115 cells closed the wound much faster than Casp^−ve^ cells. Moreover, these findings were very similar to those obtained with WM852 and WM793 cells (Figure 4G, H and S4E, F). To further validate our findings, we used a modified IncuCyte-based invasion assay in which we quantified in real time the capacity of cancer cells to migrate through a 3D matrigel plug in order to close the wound (Figure 4I for the experimental setup). As observed in the migration assays, Casp^+ve^ WM115, WM852 and WM793 cells once again displayed a greater invasive capacity (Figure 4J-M and S4G, H). Even though melanoma cells surviving apoptosis exhibited a stronger basal level of effector caspase activation (Figure S1H), the use of the pan-caspase inhibitor Q-VD-OPh prevented neither their migration nor invasion, indicating that other cell motility mechanisms must be activated (Figure S4I, J). To further validate our observations, we also used a cellular impedance-based migration assay that confirmed the enhanced migratory capacity of Casp^+ve^ WM852 cells (Figure S4K). Our findings therefore suggest that failure to undergo proper apoptosis might boost the invasive capacities of melanoma cancer cells.

### Non-lethal doses of chemotherapy enhance cancer cell aggressiveness

Dacarbazine is a cytotoxic DNA alkylating agent that has been used extensively for treatment of advanced-stage melanoma ^19^ ^20^. Although of limited effect on the overall survival, dacarbazine remains the standard treatment in countries with no routine access to BRAF-mutant or MEK inhibitors. Considering that chemotherapy is still widely used, we investigated whether dacarbazine can also trigger failed apoptosis in melanoma cells. For this, we tested a range of dacarbazine concentrations to treat WM852 or WM793 cells expressing the VC3AI caspase reporter. As depicted in the FACS plots in Figure 5A and S5A, dacarbazine treatment triggered the emergence of a VC3AI^+ve^/AnnexinV^−ve^ population both in WM852 and WM793 cells, respectively. The extent of chemotherapy-triggered failed apoptosis is quantified in Figure 5B and S5B and for subsequent experiments we used dacarbazine at a dose of 200 μg/mL. Consistent with the results obtained for tBID expression, WM852 and WM793 cells undergoing failed apoptosis following dacarbazine treatment achieve an important *in vitro* capacity to migrate and invade in both wound healing and invasion assays (Figure 5C-F and S5C-F). Although it would be interesting to extend our observations to other drugs used for melanoma treatment, these data suggest that chemotherapeutic agents could also actively favor the emergence of cancer cells failing to undergo complete apoptosis, which are much more aggressive.

**Figure 5.**
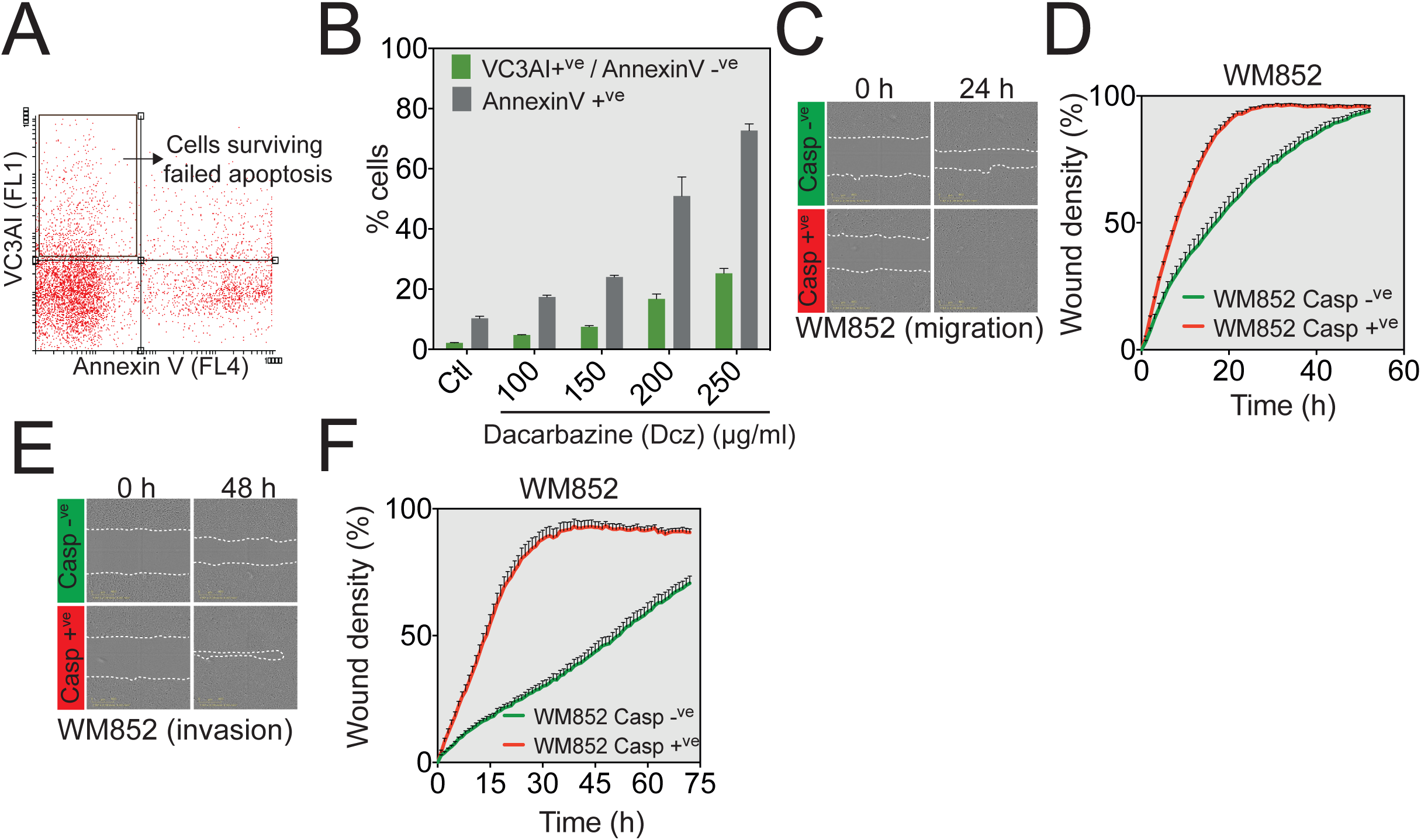
Chemotherapy can trigger failed apoptosis while increasing the invasiveness of melanoma cells. **A**. Gating strategy used to sort and further characterize WM852 cells undergoing dacarbazine (Dcz)-induced failed apoptosis (VC3AI^+ve^/AnnexinV^−ve^). **B**. WM852 cells were first treated with different concentrations of dacarbazine for 48 hours. Cells were then stained with AnnexinV-Alexa647 and analyzed by flow cytometry according to their VC3AI (FL1) expression. **C**. Monolayer WM852 cells treated with Dcz and sorted for VC3AI^+ve^/AnnexinV^−ve^ markers were wounded and photographs were taken immediately after wound induction and 24 hours later (migration assay). **D**. Measurement of the migratory capacity of WM852 through the analysis of wound area recovery using the IncuCyte imaging software. **E**. Representative images of the wound in a layer of WM852 having displayed caspase activation or not after 200 µg/mL dacarbazine treatment (invasion assay). **F**. Quantification of the invasive capacity of WM852 cells through the analysis of wound area recovery using the IncuCyte imaging software.

### Melanoma cells undergoing failed apoptosis are also more invasive *in vivo*

Given the clear gain in migratory and invasive properties displayed *in vitro* by melanoma cancer cells failing to undergo complete apoptosis, we were interested in validating our main findings *in vivo*. In recent years, the zebrafish (*Danio rerio*) has emerged as a suitable model to evaluate human cancer cell invasion within a functional circulatory system in a matter of days ^21^ ^22^. Briefly, Casp^−ve^ and Casp^+ve^ cells stained with the lipophilic dyes DiD and DiO, respectively, were injected together into zebrafish embryos, in a competition-like scenario, closer to the conditions encountered by a tumor under chemotherapy treatment (Figure 6A). Figure 6B shows representative images of zebrafish embryos displaying melanoma cells both at the site of injection in the yolk sac and throughout the tail, 24 hours after the xenograft was done. In Figure S6A the efficacy of the lipophilic staining is assessed. Mirroring the *in vitro* assays, Casp^+ve^ cells displayed a greater invasive potential compared to control cells (Figure 6C). However, when quantifying the invaded distance, there was no difference between the two populations (Figure 6D). Importantly, the invaded cells were neither fragmented nor dead, since they were stained with vital Hoechst (Figure S6B).

**Figure 6.**
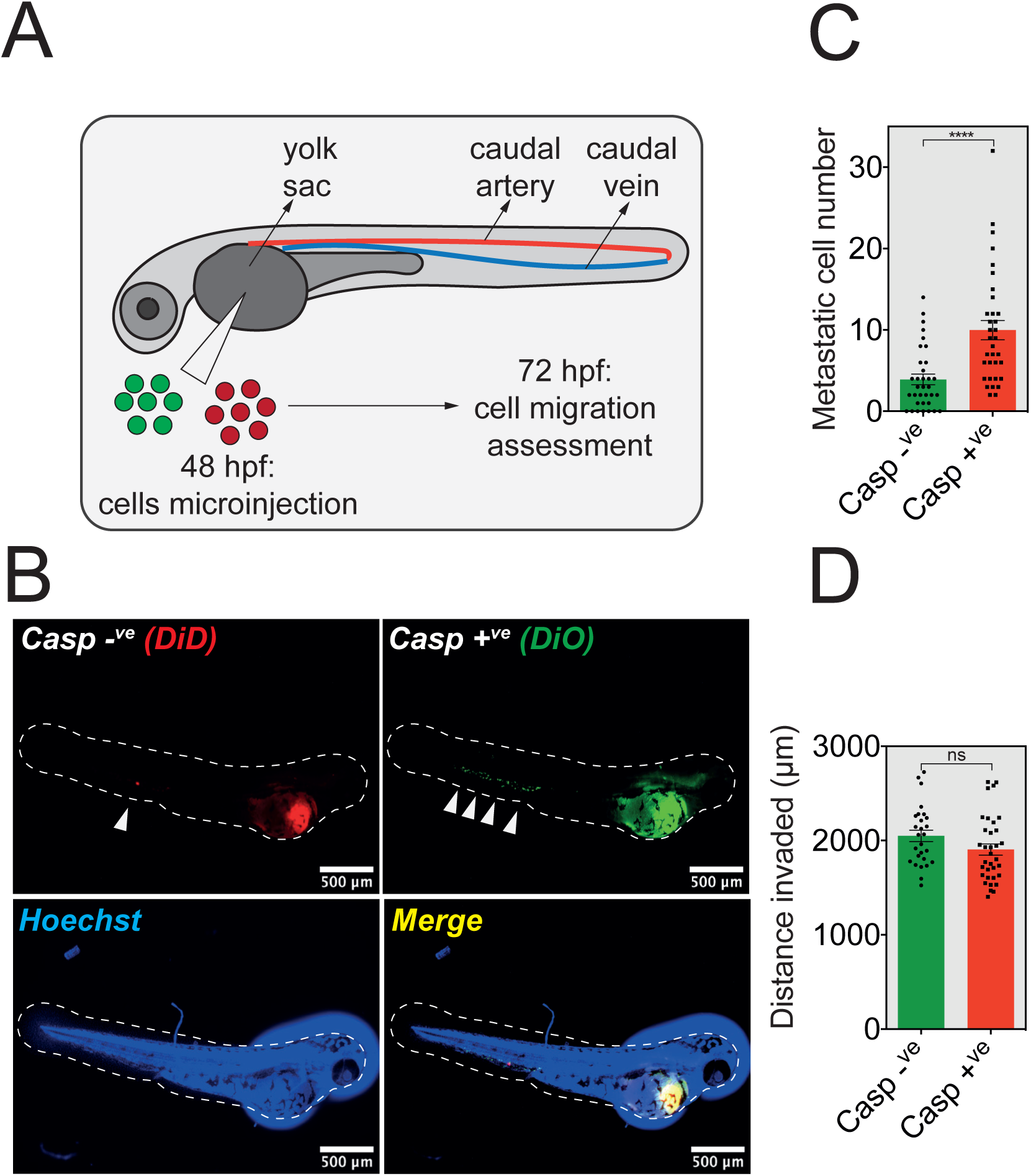
The zebrafish model of metastasis confirms the increased invasiveness of cancer cells triggered by failed apoptosis. **A.** Graphical summary of the zebrafish model of cancer metastasis. Briefly, melanoma cancer cells labeled with a lipophilic dye are injected into the perivitelline cavity of 2-day old zebrafish embryos (post-fertilization, hpf), and imaged by fluorescence microscopy 24 hours later. **B** DiD-labeled Casp^−ve^ and DiO-labeled Casp^+ve^ WM852 cells were pre-mixed in equal number and injected as described in **A**. A representative epifluorescence image of a whole embryo counter-stained with Hoechst shows both perivitelline homing and caudal blood vessel invasion of cancer cells. **C-D**. Quantification of the invaded metastatic cell number per embryo (**C**) and distance invaded from the vitellus (**D**).

To exclude a possible paracrine inhibitory effect that one cell population might exert on the other one, WM852 Casp^−ve^ and Casp^+ve^ cells stained with the lipophilic dye DiI were injected separately into the perivitelline cavity and the embryos were then analyzed for the presence of invading cells in the vasculature 24 hours post-injection (Figure S6C and S6D show representative images of zebrafish embryos displaying melanoma cells both at the site of injection in the yolk sac and throughout the tail). As presented in Figure S6E-G, the incidence of metastasis and the number of metastatic cells was higher for Casp^+ve^ cells. Collectively, these data clearly demonstrate *in vivo* the robust metastatic potential triggered by failed apoptosis.

### The JNK-AP1 transcriptional axis mediates failed apoptosis-driven cancer cell aggressiveness

Given the obvious pro-migration transcriptional signature of cells having failed apoptosis, we next wondered which transcriptional pathway might be involved in this phenotype. Previous studies showed that failure to properly execute apoptosis results in caspase-dependent DNA damage that is signaled via c-Jun N-terminal Kinase (JNK) ^7^ ^23^. This is in line with our HOMER-based motif analysis of promoter regions of all genes up-regulated following the induction of failed apoptosis in WM852 cells that revealed an enrichment in members of the AP-1 transcription factor family, such as ATF-3, Fra-1, Fra-2, BATF or JunB (Figure 7A). Since the transcriptional activity of AP-1 can be regulated upstream by JNK, we performed immunofluorescence and immunoblotting for phospho-JNK and phospho-cJun, and found that WM852 cells surviving apoptosis activate the JNK signaling pathway (Figure 7B and C). This was not the case for YAP, another pathway involved in melanoma invasion (Figure S7A). However, we were unable to detect JNK activation in unsorted WM852 cells treated with non-lethal doses of doxycycline, presumably due to the very low proportion of cells undergoing failed apoptosis (Figure S7B). To further test the involvement of JNK in driving the migration of Casp^+ve^ cells, we next treated the Casp^−ve^ and Casp^+ve^ WM852 cells with SP600125, which is a potent and selective JNK inhibitor. As expected, SP600125 reduced the migration of melanoma cells (Figure 7D and Figure S7C), and the involvement of JNK was further confirmed following JNK1/2 siRNA knockdown (Figure 7E, F). Moreover, JNK1/2 knockdown also reduces the expression of several cell motility-related genes both in WM852 and WM115 cells (Figure S7D-E).

**Figure 7.**
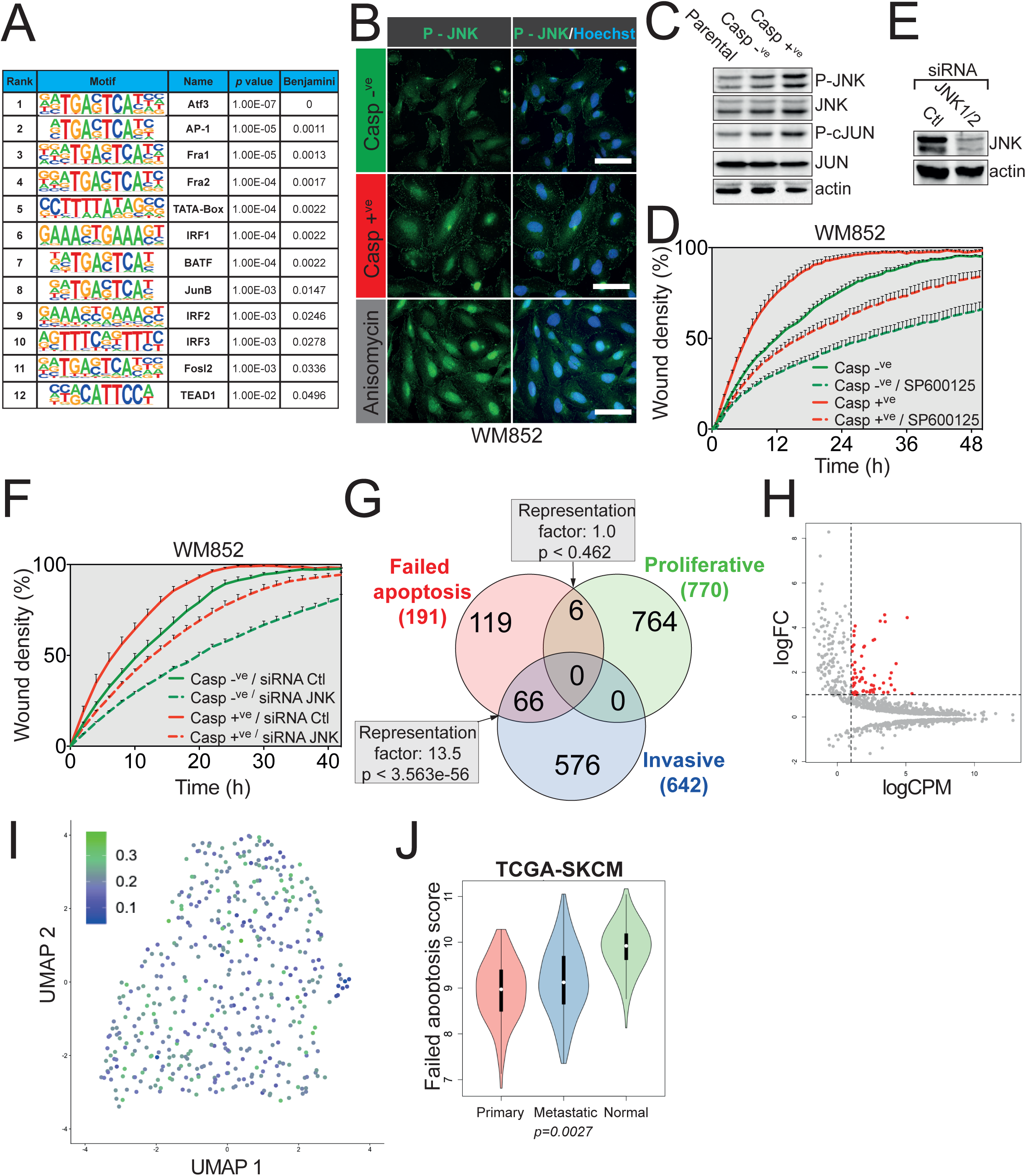
The JNK-AP1 transcriptional axis controls failed apoptosis-driven melanoma cancer invasiveness. **A.** HOMER-based motif analysis showing a significant enrichment in several transcription factors motifs in Casp^+ve^ WM852 cells. **B**. Immunofluorescence analysis for JNK phosphorylation in Casp^−ve^ and Casp^+ve^ WM852 cells. Anisomycin (25 μg/mL treatment for 1 hour) was used as a positive control. **C**. Western blot analysis of phospho-JNK and phosphor-cJUN in parental, Casp^−ve^ or Casp^+ve^ WM852 cells. Actin was used as loading control. **D**. IncuCyte ZOOM-based wound healing migration assay was performed on Casp^−ve^ or Casp^+ve^ WM852 cells treated or not with the JNK inhibitor SP600125 (10 μM). **E**. Efficacy of JNK1/2 siRNA (25 μM for each JNK1 and 2)-mediated knockdown in WM852 cells. **F**. IncuCyte ZOOM-based wound healing migration assay was performed on Casp^−ve^ or Casp^+ve^ WM852 cells transfected or not prior to the assay with siRNA control or JNK1/2 siRNA. **G**. Venn diagram detailing overlaps between the 191-genes failed apoptosis signature in WM852, the proliferative and invasive Verfaillie melanoma signature. **H**. Selection of a 60-genes signature based on their average expression, in addition to fold change and *p* value. **I**. Projection plot of the failed apoptosis score in each of the 464 malignant cells of a metastatic melanoma tumor. **J**. Comparison of the failed apoptosis score between normal skin, primary and metastatic melanoma.

Overall, these results identify the JNK signaling pathway as the top player in the migratory and invasive effects observed in melanoma cells surviving apoptosis.

We next sought to compare the changes in the transcriptome of melanoma cells experiencing failed apoptosis with published gene expression signatures. For this we selected for both WM852 and WM115 Casp^+ve^ cells a signature constituted of genes with a RNA seq expression above two fold change (191 for WM852 and 22 for WM115). As the Venn diagram in Figure 7G shows, the failed apoptosis signature in WM852 cells significantly overlaps with the “invasive” gene expression signature in melanoma, published by Verfaillie and colleagues (*p* value <3.563e-56, with 13.5 times more represented than expected by chance), while minimal overlap is observed with the “proliferative” group ^24^. The WM115 cells follow a similar trend (Figure S7F). Interestingly, failed apoptosis gene signature is different from the one published for anastasis (Figure S7G) suggesting that failed apoptosis is distinct from anastasis ^25^. To calculate a score of failed apoptosis, we further selected those genes with a higher level of expression (logCPM [counts per million] > 1), to ensure an overall higher detection across different datasets (Figure 7H). We first tested this failed apoptosis score composed of 60 genes in single-cell RNA-seq (scRNA-seq) data from a metastatic melanoma tumour sample containing 464 malignant cells ^26^. As shown in the projection plot Figure 7I, a high failed apoptosis score is detected in most metastatic melanoma cells. However, in a deconvolution analysis, we notice that the expression of several genes from the signature is highly variable among genes (Figure S7H). The analysis of publicly available TCGA data for skin cutaneous melanoma demonstrates that this signature can distinguish between primary and metastatic melanoma (Figure 7J). Intriguingly, normal skin has also a high failed apoptosis score, presumably due to an accelerated rate of cell renewal and apoptosis that might then generate statistically more failed apoptosis events. In summary, these data show that failed apoptosis gene signature might be a prognosis indicator for melanoma metastasis.

## Discussion

The main goal of this study was to determine the phenotypic consequences of failed apoptosis occurring when the apoptotic stimuli (the rising levels of pro-apoptotic proteins or chemotherapy) do not reach a lethal threshold. Even though non-lethal caspase activation has been extensively described to play an important role in differentiation, proliferation, cell migration, wound healing or iPSC reprogramming, the impact of failed apoptosis in the setting of an already established tumor is poorly described ^27^

Our data show that a partial apoptotic stimulus could be induced *in vitro* to generate a percentage of cells failing to undergo complete apoptosis. Indeed, chemotherapy does not reach and kill all tumor cells. In breast cancer, automated quantification of apoptosis one and two days after doxorubicin and docetaxel treatment revealed striking heterogeneous induction of cell death, with percentages of tumor cell apoptosis ranging from close to zero to over 35% for the same dose of chemotherapy ^28^. This is mainly due to the resilient tumor microenvironment, which inhibits the pro-apoptotic action of most cancer drugs through hypoxia, desmoplasia or inadequate tumor vascularization ^29^. Furthermore, even if the chemotherapeutic agent reaches cancer cells, its pro-apoptotic activity might be diminished by rapid metabolic changes such as cytosolic alkalinization that could mediate resistance of tumor cells to cisplatin or increased drug efflux ^30^ ^31 32^. Therefore, our failed apoptosis-induction system could also be used to mimic and better understand the effects of partial response to therapy focusing on other aspects, besides apoptosis, such as drug metabolism.

Our transcriptomic data show that a distinct gene signature centered on cell motility characterizes melanoma cancer cells undergoing failed apoptosis. Recently, the team of Denise Montell performed a whole-transcriptome RNA sequencing on anastatic cells, surviving transient apoptotic stimuli-triggered caspase activation. Interestingly, the authors described a late anastasis transcriptional program that echoes our findings since focal adhesion and regulation of actin cytoskeleton are among the top over-represented KEGG pathways. Moreover, anastatic cells stimulate cell migration through activation of TGF-β signaling and SNAIL expression, although we did not find the same specific signature possibly due to the use of different cellular models and apoptotic stimuli ^25^ ^33^. Importantly, the 60 genes-based failed apoptosis score that we describe in this study might be clinically relevant since it can distinguish between primary and metastatic melanoma. In reviewing recent literature, we also found an indication of non-lethal caspase-3 involvement in the metastatic potential of colon cancer cells. More precisely, caspase-3 KO HCT 116 cells have a reduced EMT phenotype, with a marked decrease in SNAIL, SLUG and ZEB1 ^34^. Failure to execute apoptosis also seems to impact the emergence of cancer stem cell-like cells (CSCs) in breast cancer. Here the authors found that breast cancer cells surviving non-lethal doses of staurosporine gain in metastatic potential *in vivo* while some of them acquired CSCs properties ^35^.

The tumor microenvironment, by itself or combined with classical anti-cancer therapy, is extremely hostile and therefore triggers a fight or flight reaction from cancer cells, commonly known as the adaptive stress response (ASR), which enables cancer cells to rapidly adapt to stress and survive ^36^. A known example of ASR is the epithelial-mesenchymal transition (EMT), conferring drug resistance and stem-like features to the highly invasive cancer cells ^36^. Several studies link the tyrosine kinase receptor AXL with the ASR of breast cancer or melanoma cells conferring resistance to targeted therapies, while interestingly, our study highlighted AXL as transcriptionally up-regulated in melanoma cells undergoing failed apoptosis ^26, 37^. Moreover, the transcription factor ATF3, at the apex of our transcriptional regulation network characterizing failed apoptosis in melanoma, was also described as the central hub of the ASR ^36^. Therefore we could speculate that failed apoptosis is a defense mechanisms triggered by the ASR, allowing cancer cell survival and metastasis.

The key finding to emerge from this study is that melanoma cancer cells impacted by failed apoptosis gain in motility. IncuCyte-based live-cell imaging revealed that these cells display a collective migration, as cell sheets and clusters. This type of migration characterizes several types of cancers, such as prostate, large cell lung cancer, melanoma or rhabdomyosarcoma ^38, 39^. The mechanisms underlying collective cell migration have best been studied in Drosophila, where JNK was shown to regulate border cells migration as cellular sheets ^40^. In accordance with this, our results also demonstrate that the JNK pathway appears to regulate failed apoptosis-driven melanoma migration. Our findings match those observed in earlier studies. For instance, constitutive MAPK activation frequently observed in melanoma, up-regulates JNK and activates the oncogene c-JUN by increasing its transcription and stability ^41^. In addition, the transition from proliferative to invasive melanoma was shown to be orchestrated by the AP-1/TEAD, which are the transcriptional master regulators downstream of JNK and Hippo pathways, respectively ^24^. Of note, our RNA seq data fits well with the invasive signature published by Verfaillie and colleagues, while no correlation is observed with the proliferative signature. In addition, the failed apoptosis signature can also be detected in most of cells of a metastatic melanoma tumour analyzed by scRNA-seq. Aside from melanoma, AP-1/TEAD modulates the expression of a core set of genes involved in the migration and invasion of different types of cancer cells, including neuroblastoma, colorectal or lung cancer ^42^. Regarding the AP-1 transcriptional targets it is worth noting that melanoma cells failing to die up-regulate several genes such as *PTX3*, *MYLK*, *SPP1* or *RAC2*, all shown to be regulated by AP-1 and involved in metastasis ^43^ ^44^ ^45^ ^46^. While in these studies the AP-1/c-JUN transcription factors are activated by a plethora of stimuli such as cytokines, growth factors, UV irradiation or activating mutations in N-RAS and B-RAF genes, in our case, JNK activation was most probably induced as a consequence of minority MOMP ^7^. Indeed, it was shown that limited mitochondrial permeabilization and subsequent non-lethal caspase activation could induce DNA damage and, as a consequence, activate JNK ^2,7^. In our setup, however, continuous or pulsed treatment during migration assays with the pan-caspase inhibitor Q-VD-OPh did not prevent failed apoptosis-induced migration, indicating that either caspases do not play a role in these settings or they act punctually to kick-start the migratory program and are then dispensable. Another explanation might be that failed apoptosis-induced aggressiveness is entirely MOMP-dependent and caspase-independent. Along this line, MOMP-triggered SMAC release could mediate the protein stability of Rho GTPase RAC1 *via* the repression of XIAP and cIAP1, two anti-apoptotic proteins shown to act as E3 ubiquitin ligases for RAC1 ^47^. As intriguing as it might be, further research is required to investigate this hypothesis.

Even though we used melanoma cancer cells as a proof-of-principle tool to investigate failed apoptosis, we hypothesize that our findings might extend to other types of cancers. In this vein, research conducted in colon or breast cancer cells also highlighted a correlation between increased aggressiveness and non-lethal caspase activation ^35,48^.

In conclusion, our work demonstrates that the resilience of cancer cells to undergo complete apoptosis leads to a more aggressive cancer phenotype. Moreover, we provide a model to induce and study the effects of failed apoptosis in physiological settings such as neuronal function or stemness. From a clinical perspective, elucidating the functional significance of failed apoptosis-driven transcriptional signature and underlying mechanisms may improve the targeting of metastatic processes.

## Material and methods

### Cell culture and treatments

Melanoma cell lines (WM115, WM793, WM852, WM239A, 501Mel and WM35) were provided by either Patrick Mehlen (CRCL, France) or Robert Insall (Beatson Institute, UK). Cells were grown in Dulbecco’s Modified Eagle’s Medium supplemented with 10% FCS (Eurobio, CVFSVF00-01), 2 mM glutamine (ThermoFisher Scientific, 25030-024), non-essential amino acids (ThermoFisher Scientific, 11140-035), 1 mM sodium pyruvate (ThermoFisher Scientific, 11360-039) and penicillin/streptomycin/ampicillin (ThermoFisher Scientific, 15140-122). Cells were regularly checked for mycoplasma contamination. Other reagents used in this study: Q-VD-OPh (Clinisciences, JM-1170), SP600125 (Sigma-Aldrich, S5567), doxycycline (Sigma-Aldrich, D9891), SYTOX Green (ThermoFisher Scientific, S34860), propidium iodide (Sigma-Aldrich, P4864), Hoechst 33342 (ThermoFisher Scientific, H1399) and dacarbazine (Sigma-Aldrich, D2390).

### Lentiviral transduction and cell lines generation

The plasmid encoding for the caspase reporter VC3AI, namely pCDH-puro-CMV-VC3AI, was purchase from Addgene, cat.no. 78907. For the lentiviral transduction, 293T cells (2×10^6^ in a 10 cm dish) were transfected with pCDH-puro-CMV-VC3AI using Lipofectamine 2000 (ThermoFisher Scientific, 11668019) according to the manufacturer’s instructions. The helper plasmids for lentiviral production were pVSVg (Addgene, 8454) and psPAX2 (Addgene, 12260). Two days later, virus-containing supernatant was harvested, filtered and used to infect target cells in the presence of 1 μg/mL polybrene. Two days post-infection, stably expressing cells were selected by growth in 1 μg/mL puromycin. Melanoma cells with Dox-inducible expression of tBID protein were created using the Sleeping Beauty transposon system ^49^. The plasmids that were used are pSBtet-RB (Addgene, 60506) pSBtet-BB (Addgene, 60505) and pCMV(CAT)T7-SB100 (Addgene, 34879) encoding for the SB100X transposase. First, Flag-tBID was cloned into the SfiI site of pSBtet-RB and pSBtet-BB as a PCR fragment amplified using the forward primer CGCGGCCTCTGAGGCCATGGATTACAAGGATGATGATGATAAG and the reverse CGCGGCCTGTCAGGCCTCAGTCCATCCCATTTCTGG. The pSBtet-RB-FLAGtBID and pSBtet-BB-FLAGtBID were then co-transfected with the SB100X transposase in melanoma cells using Lipofectamine 2000 and selected with blasticidin at 5 μg/ml.

### siRNA transfection

siRNAs targeting JNK1 (M-003514-04) and JNK2 (M-003505-02) were purchased from Dharmacon. 10^6^ WM852 cells were plated in 10-cm petri dishes 24 hours prior to transfection. Cells were then transfected with 25 µM siRNA using lipofectamine RNAiMAX according to the manufacturer’s instructions (ThermoFisher Scientific, 13778150).

### Cell adhesion assay

96-Well imageLock (Sartorius, 4379) plates were coated with Matrigel (Sigma Aldrich, E609-10mL) at 100 µg/mL for 1 hour at 37°C. The excess of Matrigel was then removed and 10^3^ melanoma cells (WM793 or WM852) were seeded onto the remaining thin layer. Cell images were then acquired at different time intervals using the IncuCyte ZOOM Imaging System (Sartorius). Cell adhesion was determined through cell shape analysis and scoring.

### Wound healing assay

3×10^4^ WM852 or 5×10^4^ WM115/WM793 cells were seeded in a 96-Well imageLock plate (Sartorius, 4379) and allowed to form a cell monolayer for 24 hours prior to the assay. Next, a wound was done in the cell monolayer using the WoundMaker (Sartorius, 4563), according to the manufacturer’s instructions. Cell migration to close the wound was then assessed by time-lapse microscopy using the IncuCyte ZOOM imaging system.

### Invasion assay

96-Well imageLock plate (4379, Sartorius) were coated with Matrigel (Sigma Aldrich, E609-10mL) at 100 µg/mL for 1 hour at 37°C. The excess Matrigel was removed and 3×10^4^ WM852 or 5×10^4^ WM115/WM793 cells were seeded 24 hours prior to the assay. Next, a wound was done in the cell monolayer using the WoundMaker and a new layer of Matrigel (800 µg/mL) was loaded onto cells and allowed to polymerize for 1 hour at 37°C. Medium was then added on top of the well and the invasion capacity of melanoma cells was evaluated using the IncuCyte ZOOM-based time-lapse microscopy.

### Chemotaxis assays

Melanoma chemotaxis was evaluated using 8 µm-transwells (Corning, 3422). Briefly, 10^5^ melanoma cells (WM115, WM793 or WM852) were seeded onto the upper compartment of a Boyden chamber in 300 µL of serum-free DMEM. The lower chamber was filled with 500 µL of media supplemented with 20% serum. 48 hours later, cells that had passed through the membrane pores were stained for 15 minutes with Hoechst 33342 (1 µg/mL) in PBS and counted in three representative areas by two researchers. Alternatively, melanoma chemotactic capacities were determined using xCELLigence technology. 3×10^4^ melanoma cells (WM115 or WM852) were seeded onto the upper chamber of a cim-plate 16 (ACEA Biosciences, 05665817001) and the lower chamber was filled with medium supplemented with 20% serum. Melanoma cell migration was then followed over time using the xCELLigence RTCA DP instrument (ACEA Biosciences).

### Apoptosis assay

5×10^4^ WM793 or WM852 were seeded onto 24-well plates and treated with various concentrations of doxycycline in presence of 0.3 µg/mL propidium iodide or 30 nM SYTOX Green. Cells were then imaged every 60 or 120 minutes using the IncuCyte ZOOM imager and the number of PI/SYTOX Green positive cells was normalized against the initial confluency factor of the respective well.

### RNA sequencing and quantitative RT-PCR

Total RNA was extracted using the Nucleospin RNA extraction kit (Macherey Nagel, 740955) according to the manufacturer’s instruction. For RNAseq analysis, libraries were prepared from 600 ng total RNA per sample with TruSeq Stranded mRNA kit (Illumina) following the manufacturer’s recommendations. The key steps consist of PolyA mRNA capture with oligo dT beads, cDNA double strand synthesis, and ligation of adaptors, library amplification and sequencing. Sequencing was performed using the NextSeq500 Illumina sequencer in 75 bp paired-end.

For mRNA relative expression analysis by qRT-PCR, mRNAs were first converted into cDNA using the Sensifast cDNA synthesis kit (Bioline, BIO-65053). Specific primers for each gene of interest were designed using Primer-blast website (https://www.ncbi.nlm.nih.gov/tools/primer-blast/) and are listed in Table n°1. GAPDH was used as the invariant control. The thermal cycling conditions comprised an initial polymerase activation step at 95 °C for 2 minutes, followed by 40 cycles at 95 °C, 5 seconds, and 60 °C, 30 seconds.

**Table 1.**
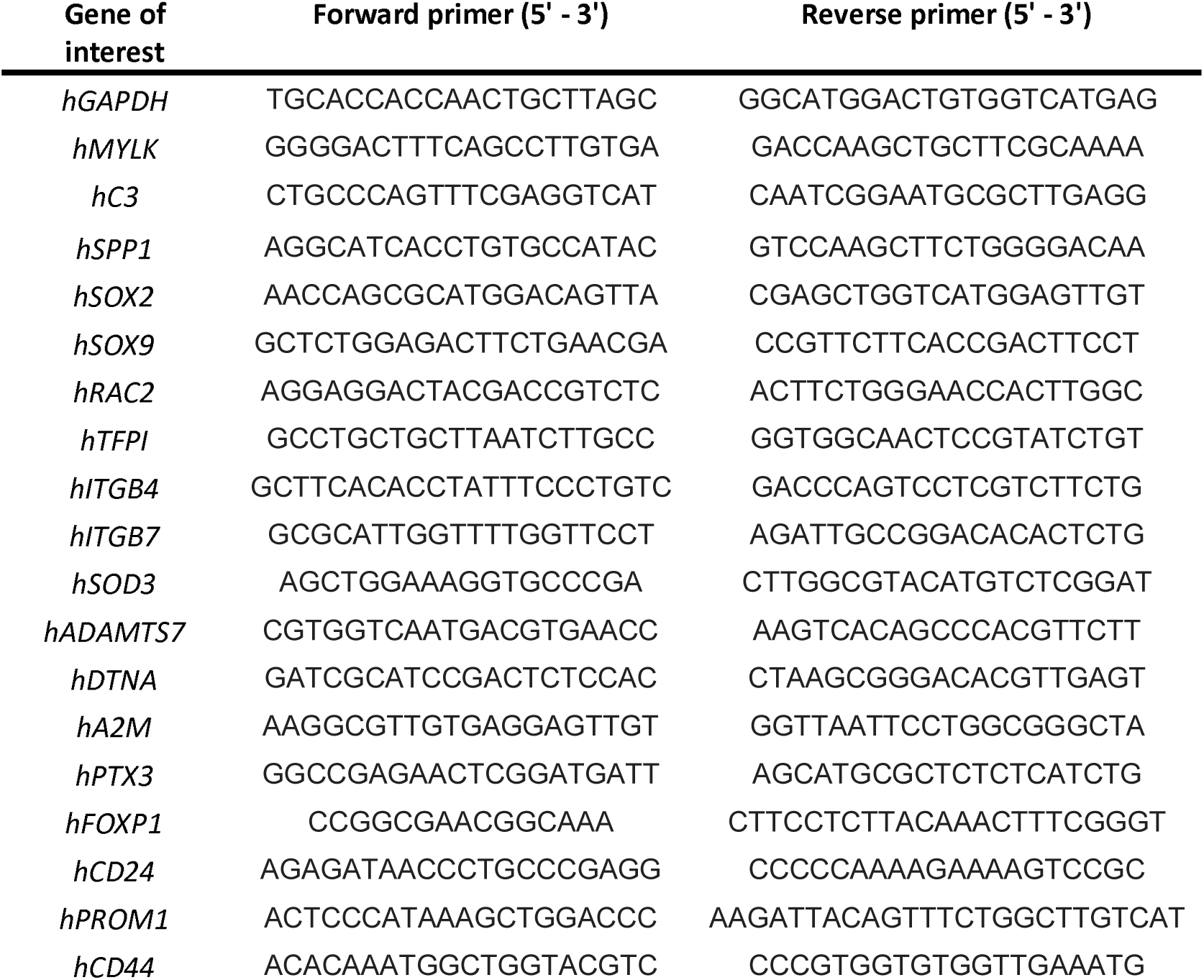
qRT-PCR primers used in this study.

### Western Blot

Protein extraction was performed using RIPA buffer (Cell signaling, 9806S) supplemented with protease inhibitor cocktail (Sigma-Aldrich, 4693116001) and phosphatase inhibitors (Sigma Aldrich, P5726-1ML, P6044-1ML). Proteins were quantified using the Protein Assay dye Reagent Concentrate (Biorad, 5000006). 50 µg of each sample were then separated in SDS-PAGE and transferred onto a nitrocellulose membrane using Transblot Turbo Transfer System (Biorad, 1704150EDU). Non-specific binding sites on the membranes were then blocked in 5% BSA or milk in TBS-Tween 0.1% for 45 minutes. Membrane were then incubated overnight at 4°C under agitation with primary antibody diluted at 1/1000 in 1% BSA TBS-tween 0.1% (FLAG (Sigma-Aldrich, F3165), P-JNK (Cell signaling, 4668), JNK (Cell signaling, 9252), P-cJun (Cell signaling, 9164), cJun (Cell signaling, 9165), cPARP (Cell signaling, 9532), actin (Sigma-Aldrich, A3854)). Membranes were then washed 3 × 10 minutes in TBS-T. The appropriate secondary antibody coupled to the horseradish peroxidase (Biorad, 1706515 and 1706516) was then added for 45 minutes at room temperature under agitation. Membranes were washed 3 times and proteins were detected using Clarity Western ECL blotting substrates (Biorad, 1705060) and chemiDoc imager (Biorad, 17001401).

### Immunofluorescence

5×10^4^ WM115, WM793 or WM852 were seeded onto coverslips eventually coated with Matrigel for focal adhesion associated protein analysis, in 24-well plates for 24 hours. Cells were then fixed in 4% PFA for 5 minutes and washed in PBS. Cells were permeabilized using 0.2% Triton X-100 in PBS for 10 minutes and non-specific binding sites were blocked using 2% BSA in PBS. Cells were incubated with the primary antibody ((FLAG (1/1000, Sigma-Aldrich, F3165), Paxillin (1/400, BD Transduction Biosciences, 610052), Vinculin (1/500, Sigma-Aldrich, V9131)) for 1 hour at room temperature or overnight in the cold room. After three washes, cells were incubated with the appropriate secondary antibody coupled to Alexa Fluor (1/300, Thermofisher scientific, A21151 and A31571) for 1 hour at room temperature protected from light. Nuclei were stained with Hoechst 33342 (10 µg/mL, ThermoFisher Scientific, H1399). Cells were washed and coverslips were mounted using fluoromount (Southern Biotech, 0100-01) before image acquisition using a Zeiss Axio Imager (Zeiss).

### Cell sorting using flow cytometry

Melanoma cell lines were treated for 24 hours with doxycycline (100 ng/mL for WM852 and 250 ng/mL for WM115 and WM793) or 48 hours with 200 µg/mL of dacarbazine. All cells were harvested, stained with AnnexinV-Alexa Fluor 647 (Biolegend, 640912) and 1 µg/mL DAPI (Sigma-Aldrich, D9542) according to the manufacturer’s instructions. Cells positive for caspase activation and negative for AnnexinV and DAPI were sorted on a FACS ARIA (BD Biosciences).

### Caspase 3 fluorometric assay

Caspase 3 activity was determined using the Caspase 3/CPP32 Fluorometric assay kit according to the manufacturer’s instructions (BioVision, K105)

### Zebrafish metastasis model

Prior to injection, 9×10^5^ melanoma cells were resuspended in serum-free medium and stained with lipophilic dyes DiO or DiD for 20 minutes at 37°C (ThermoFisher Scientific, V22889). Cells were then washed and resuspended in 30 µL of PBS. For zebrafish xenotransplantation, 48 hours post-fecundation (hpf) zebrafish embryos were dechorionated and anaesthetized with tricaine (Sigma-Aldrich, E10521) and 20 nL of cell suspension (approximately 300 labeled human cells) were injected into the perivitelline cavity of each embryo. The embryos were then placed at 30°C for 24 hours and allowed to recover in the presence of N-phenylthiourea (Sigma-Aldrich, P7629) to inhibit melanocyte formation. For imaging and metastasis assessment, zebrafish embryos were anaesthetized with tricaine and imaged using an Axio Observer Zeiss microscope (Zeiss).

### Bioinformatic analyses

All genomic data was analysed with R/Bioconductor packages, R version 3.6.1 (2019-07-05) [https://cran.r-project.org/; http://www.bioconductor.org/].

#### RNA-Seq differential analysis

Illumina sequencing was performed on RNA extracted from triplicates of each condition. Standard Illumina bioinformatics analyses were used to generate fastq files, followed by quality assessment [MultiQC v1.7 https://multiqc.info/], trimming and demultiplexing. ‘Rsubread’ v1.34.6 was used for mapping to the hg38 genome and creating a matrix of RNA-Seq counts. Next, a DGElist object was created with the ‘edgeR’ package v3.26.7 [https://doi.org/10.1093/bioinformatics/btp616]. After normalization for composition bias, genewise exact tests were computed for differences in the means between groups, and differentially expressed genes (DEGs) were extracted based on an FDR-adjusted p value < 0.05 and a minimum absolute fold change of 2. DEG gene symbols were tested for the overlap with published signatures of interest using a hypergeometric test. Hypergeometric Optimization of Motif EnRichment (HOMER v3.12) [https://doi.org/10.1016/j.molcel.2010.05.004] was used to calculate motif enrichment on the promoters of DEGs (up- and down-regulated genes separately), using default background settings.

#### Failed apoptosis score

To calculate a score of failed apoptosis, we selected DEGs based on their average expression (logCPM > 1), in addition to their fold change (logFC > 1) and p value (FDR < 0.05). A score was created using the sig.score function of the ‘genefu’ package v.2.11.2 [https://doi.org/10.1093/jnci/djr545], with logarithmic fold-change (logFC) as coefficient giving the direction and strength of the association of each gene in the gene list. This score was applied to single cell data (see below), primary and metastatic melanoma gene expression data from TCGA (SKCM), and normal skin expression data from the Genotype-Tissue Expression (GTEx) repository [https://gtexportal.org/home/].

#### Single Cell RNA-Seq Reanalysis

Single cell data was analysed with the ‘Seurat’ package v.3.1.0 [https://doi.org/10.1016/j.cell.2019.05.031]. Published metastatic melanoma single cell RNA-Seq data [https://doi.org/10.1126/science.aad0501] was downloaded from the GEO repository [https://www.ncbi.nlm.nih.gov/geo/query/acc.cgi?acc=GSE72056] as normalized expression levels. To account for inter-individual variation, we selected malignant cells from the subject with the highest number of cells (n= 464 malignant cells). Then, a Seurat object was created, normalized, and re-scaled on this subset. A failed apoptosis score was calculated for each melanoma cell, as described above, and plotted using the Uniform Manifold Approximation and Projection (UMAP) dimensional reduction technique [https://arxiv.org/abs/1802.03426].

#### TCGA Data

Melanoma cancer expression and clinical data was downloaded from the The Cancer Genome Atlas (TCGA) repository [https://www.cancer.gov/tcga.] using the R package ‘TCGAbiolinks’ v.2.9.4 [http://doi.org/10.1093/nar/gkv1507]. Briefly, TCGA-SKCM was queried for Illumina HiSeq RNA-Seq normalized results (n=471 samples). Downloaded data was pre-processed and a failed apoptosis score was calculated for each sample, as described above. Failed apoptosis scores were then compared in Primary (n=103) vs. Metastatic (n=368) melanoma groups.

## Statistical analysis

For comparison of multiple groups, two-way Analysis of Variance (ANOVA) was used while Student’s t test was applied when comparing two groups. Analyses were performed using Prism 5.0 software (GraphPad).

## Acknowledgements

This work was supported by funding from LabEX DEVweCAN (University of Lyon), Fondation ARC pour la recherche sur le cancer (grant n° 20171206348), Agence Nationale de la Recherche (ANR) Young Researchers Project (ANR-18-CE13-0005-01) and the ANR PLAsCAN Institute. We thank Brigitte Manship for reviewing the manuscript, Florine Mugnier and Sara Guedda for technical assistance and David Bernard for discussing this project.

## Figure legends

**Supplementary Figure 1.**
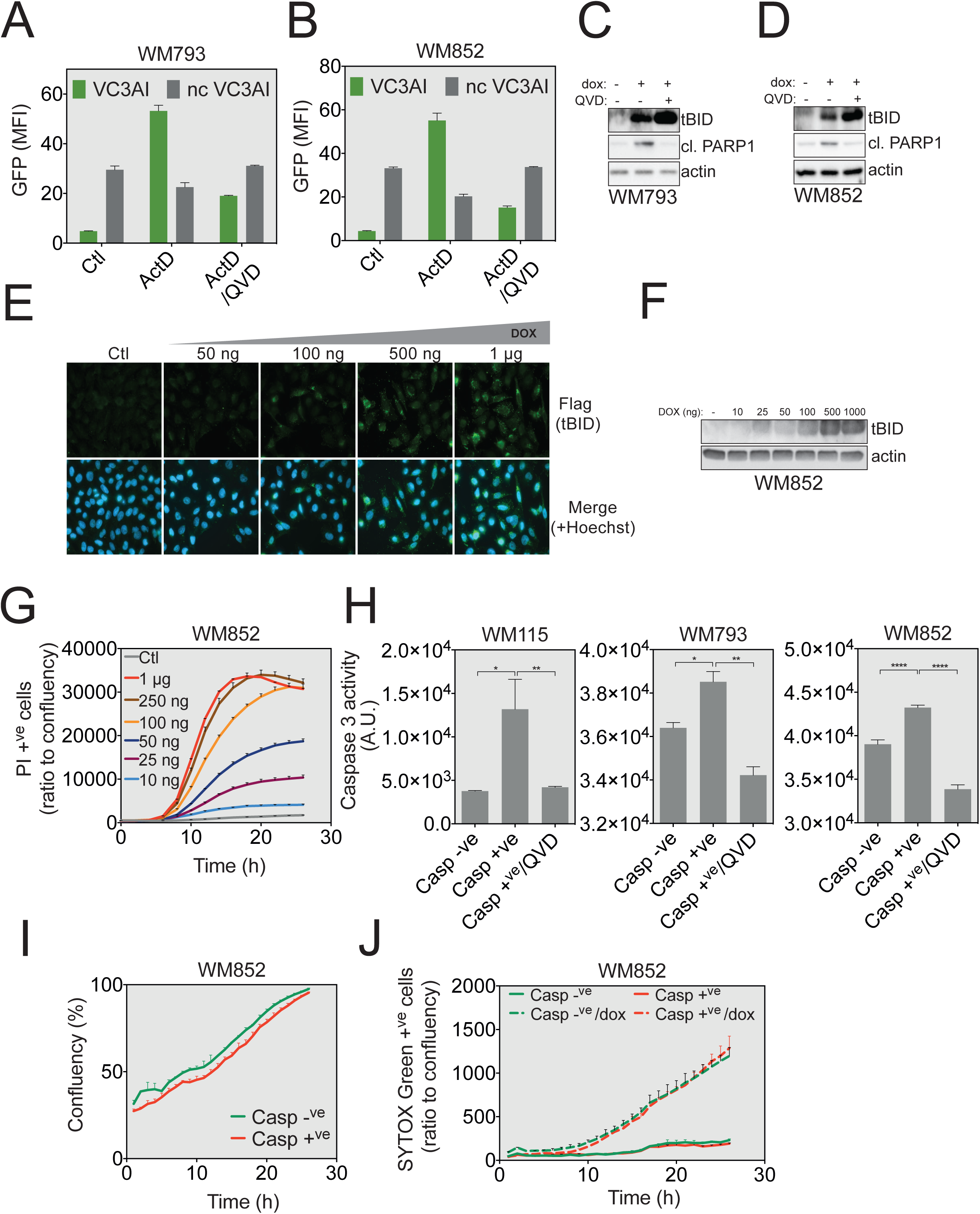
Related to Figure 1. **A**. WM793 cells expressing both the VC3AI reporter and tetON tBID system, were treated with 1 µM actinomycin D in the presence or absence of 10 µM QVD for 24 hours. Caspase activation was then determined by flow cytometry. Cells stably expressing a non-cleavable form of VC3AI (ncVC3AI) served as control. **B**. WM852 cells were treated and analyzed as described in **A**. **C-D**. Immunoblot analysis of tBID expression and cleaved PARP1 in WM793 (**C**) and WM852 cells (**D**) treated with 1 µg/mL of doxycycline with or without QVD for 24 hours. Actin served as a loading control. **E**. WM115 cells were treated with various concentrations of doxycycline while tBID expression was determined by immunofluorescence using an anti-Flag immunostaining. Hoechst was used for nuclear staining. **F**. WM852 cells were treated with various concentrations of doxycycline for 24 hours and tBID expression was determined by Western blot. Actin served as a loading control. **G**. WM852 cells were treated with various concentrations of doxycycline and apoptosis induction was measured in real-time using PI staining and IncuCyte ZOOM-based live-cell imaging. **H**. Effector caspase activation in sorted cells (7 days post-sorting) was measured in the presence or absence of QVD for 24 hours using the Caspase-3 fluorometric Assay kit. **I**. The proliferation of WM852 cells failing to undergo apoptosis (the Caspase +^ve^ sorted population) was determined using the IncuCyte ZOOM live-cell imaging based on measuring the confluency parameter. **J**. Responsiveness to a new tBID expression following doxycycline treatment was measured using Sytox staining and the Incucyte Zoom Imaging System. **J**. The responsiveness of WM852 cells to a new tBID-expression challenge was measured using SYTOX Green staining.

**Supplementary Figure 2.**
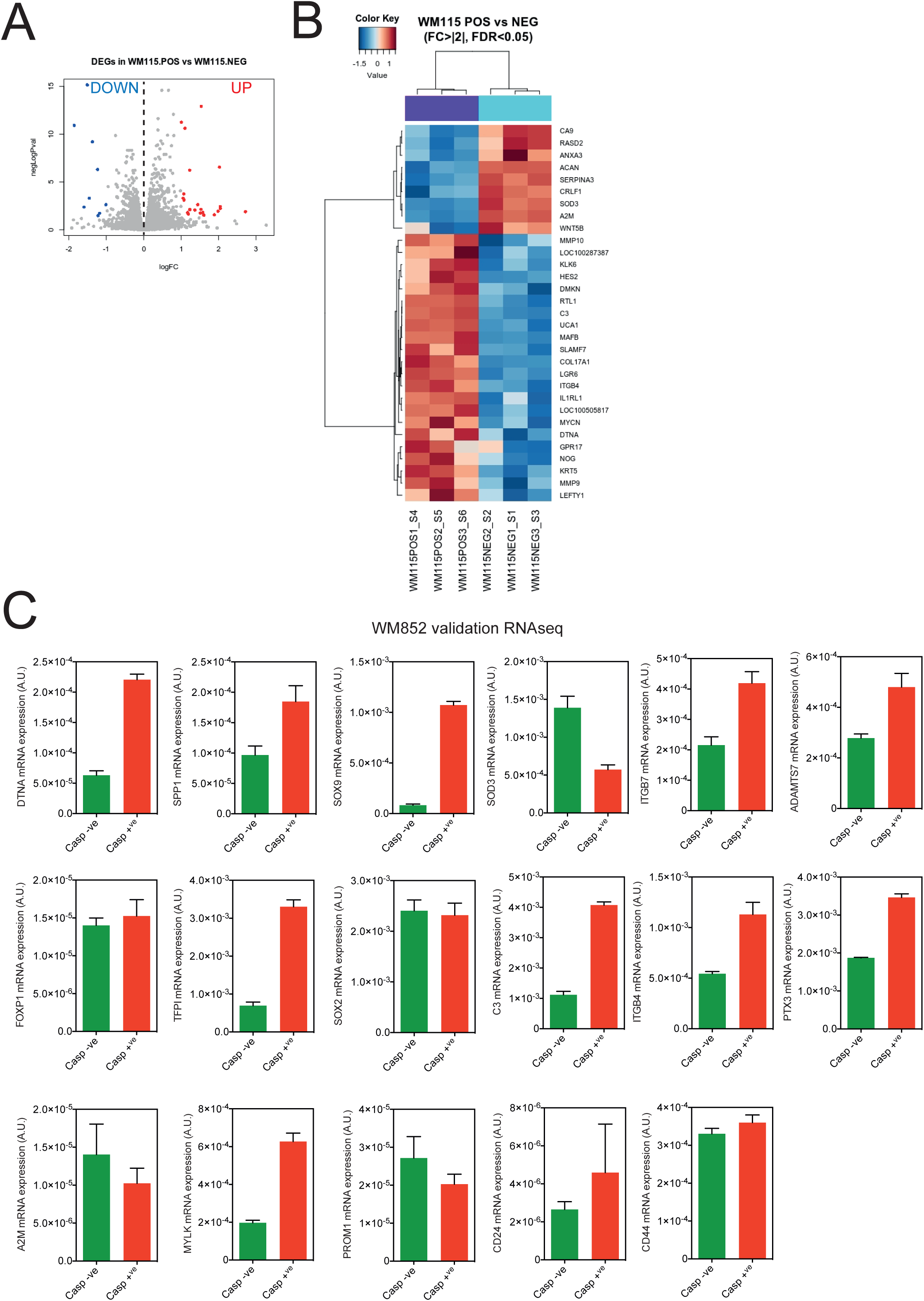
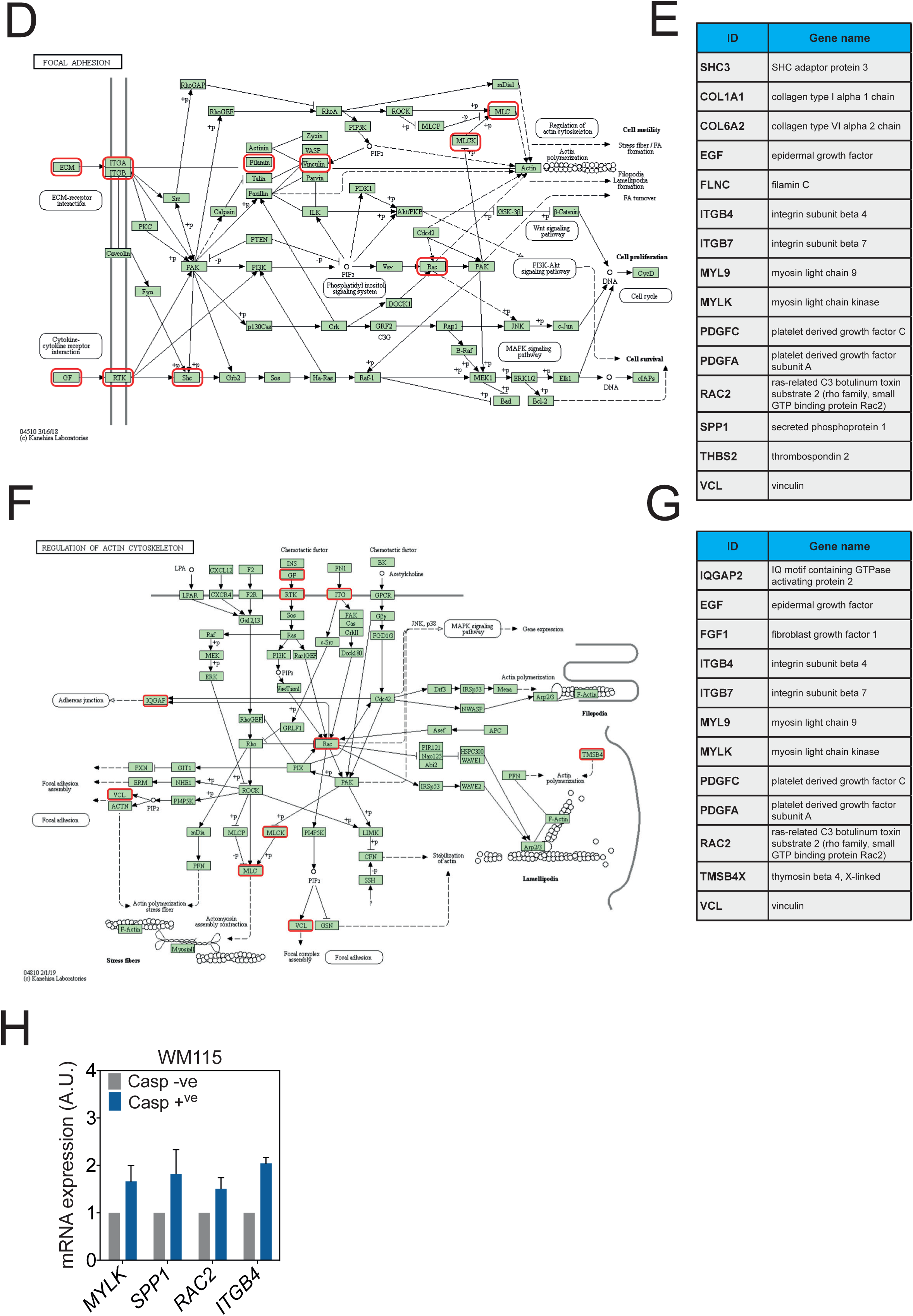
Related to Figure 2. **A**. Vulcanoplot displaying the expression (in log fold change) of each gene differentially expressed in WM115 Casp^+ve^ compared to WM852 Casp^−ve^ cells. **B**. Unsupervised clustering of the RNAseq data in WM115 Casp^−ve^ and Casp^+ve^ cells. Red indicates increased while blue indicates decreased mRNA abundance of selected genes with fold change above 2. **C**. Random selection of 17 genes used for RNAseq validation in WM852 by q-RT-PCR. **D**. “Focal adhesion” KEGG pathway. Genes identified in WM852 RNAseq are highlighted with a red frame. **E**. The list of genes identified in WM852 cells undergoing failed apoptosis, associated with the “Focal adhesion” KEGG pathway. **F**. “Regulation of actin cytoskeleton” KEGG pathway. Genes identified in WM852 RNAseq are highlighted with a red frame. **G**. The list of genes identified in WM852 cells sustaining failed apoptosis, associated with “Regulation of actin cytoskeleton” KEGG pathway. **H**. q-RT-PCR on WM115 Casp^−ve^ and Casp^+ve^ cells for selected genes involved in cell motility, common with WM852 cells.

**Supplementary Figure 3.**
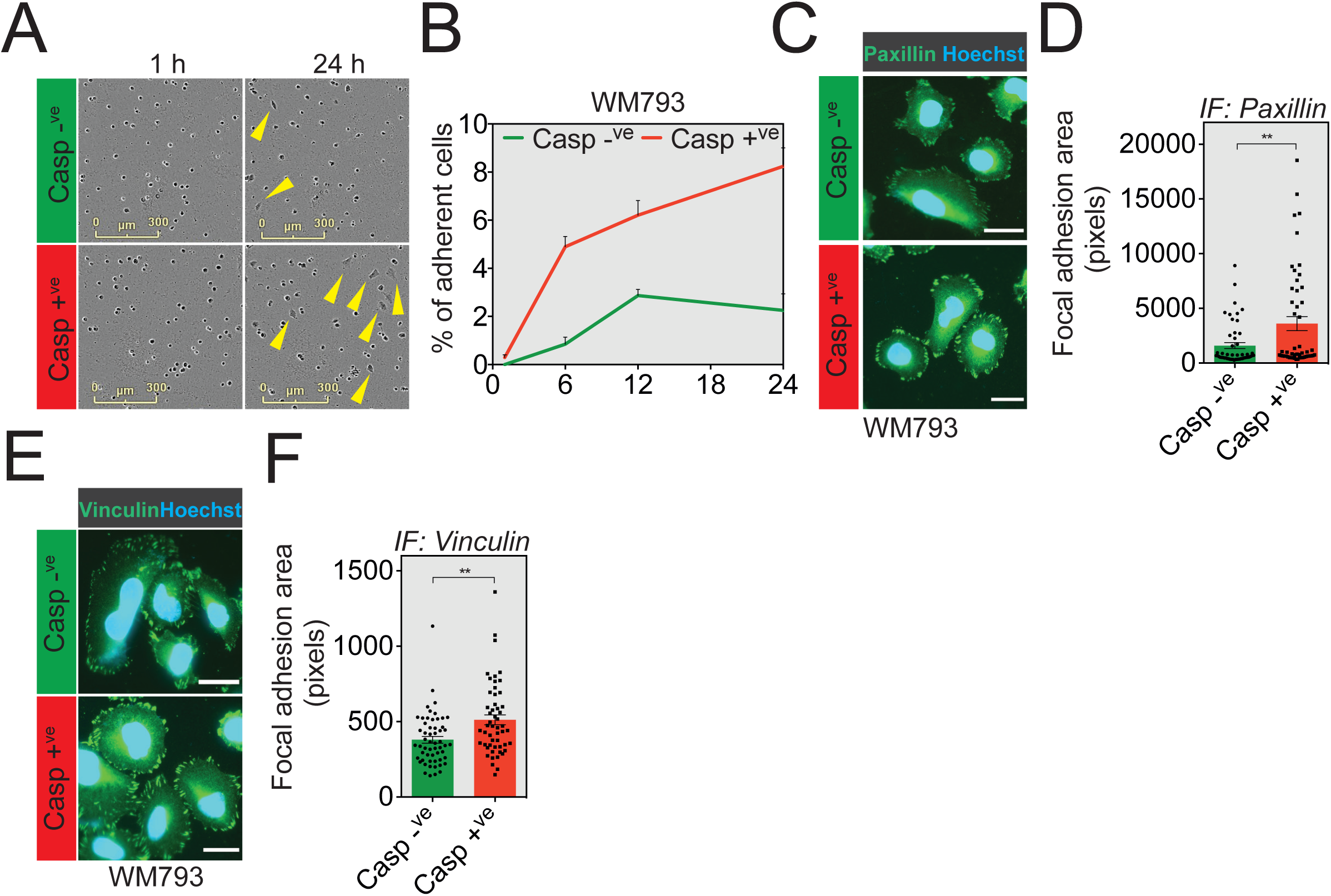
Related to Figure 3. **A**. Casp –^ve^ and Casp +^ve^ WM793 melanoma cells were seeded onto a 96-well plate previously coated with 100 µg/mL matrigel and imaged using the IncuCyte ZOOM. Representative images of WM793 adhesion capacities are shown at 1h and 24 hours after seeding. **B**. Quantification of the number of WM793 adherent cells. **C**. Paxillin immunostaining analysis in Casp^−ve^ and Casp^+ve^ sorted WM793 melanoma cells. **D**. Quantification of focal adhesion area based on paxillin immunostaining. **E**. Vinculin immunostaining analysis in Casp^−ve^ and Casp^+ve^ sorted WM793 melanoma cells. **F**. Quantification of focal adhesion area based on vinculin immunostaining.

**Supplementary Figure 4.**
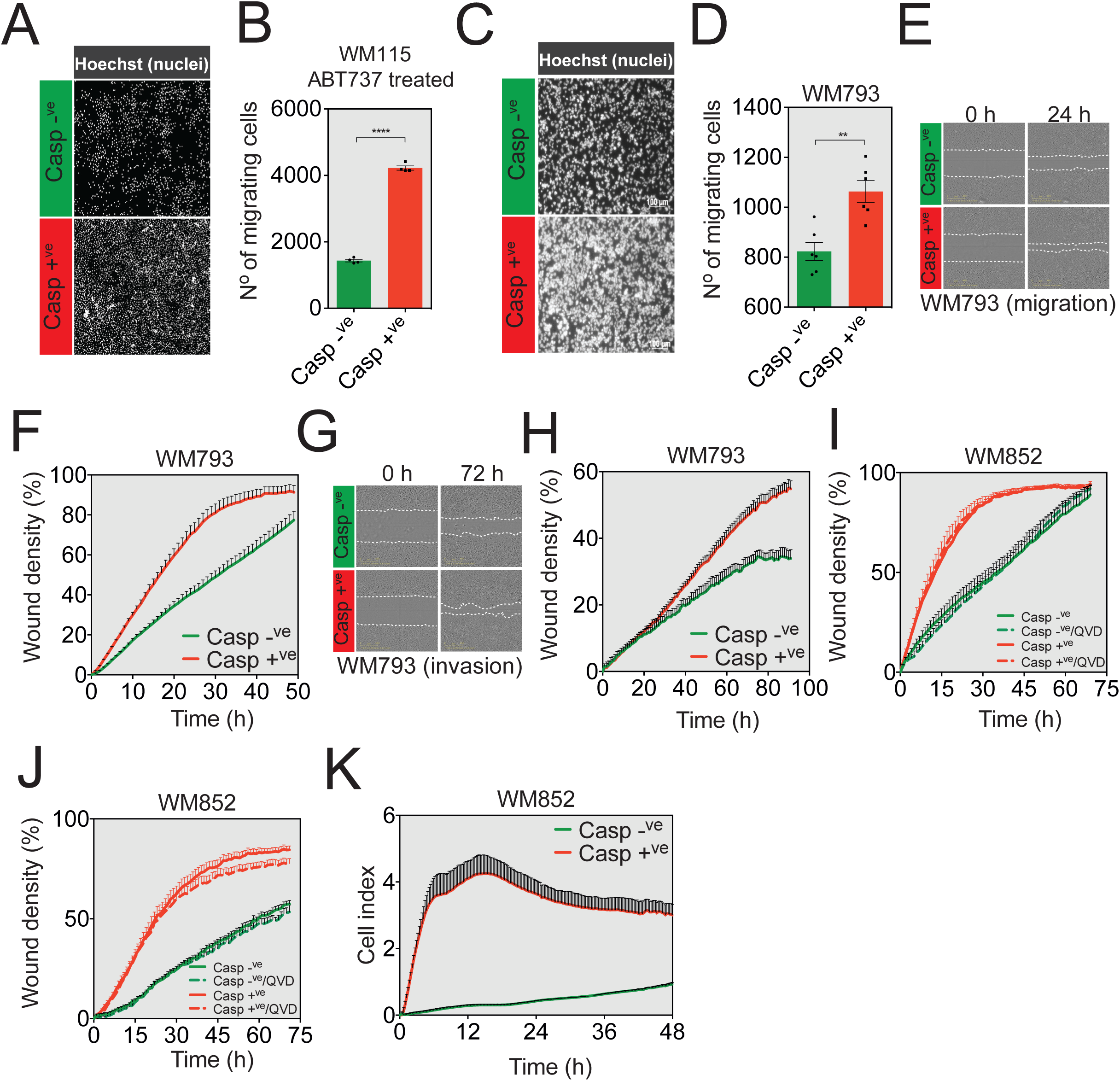
Related to Figure 4. **A**. Representatives images of chemotactic potential of Casp^−ve^ and Casp^+ve^ sorted WM115 following their treatment with the BH3 mimetic ABT-737. **B**. Quantification of the number of WM115 cells undergoing ABT-737-triggered failed apoptosis having crossed the transwell membrane. **C**. Representative images of Hoechst-stained Casp^−ve^ and Casp^+ve^ sorted WM793 cells that passed through a transwell membrane. **D**. Quantification of the number of WM793 having migrated through an 8-µm transwell membrane. **E**. Monolayer WM793 cells were wounded and photographs were taken immediately after wound induction and 24 hours later. **F**. IncuCyte ZOOM-based quantification of the migratory potential of sorted WM793 cells through wound area measurement. **G**. Representative images of the wound in a layer of sorted WM793 cells at 0 and 72 hours after wounding in the presence of matrigel. **H**. IncuCyte ZOOM-based measurement of WM793 cell invasion through matrigel. **I**. Effect of caspase inhibition following Q-VD-QPh (10 µM) treatment on the migratory capacity of WM852 cells after sorting for caspase activation. **J**. Same is in (I) while the quantification is performed to assess the invasive potential. **K**. xCELLigence cellular impedance chemotaxis assay.

**Supplementary Figure 5.**
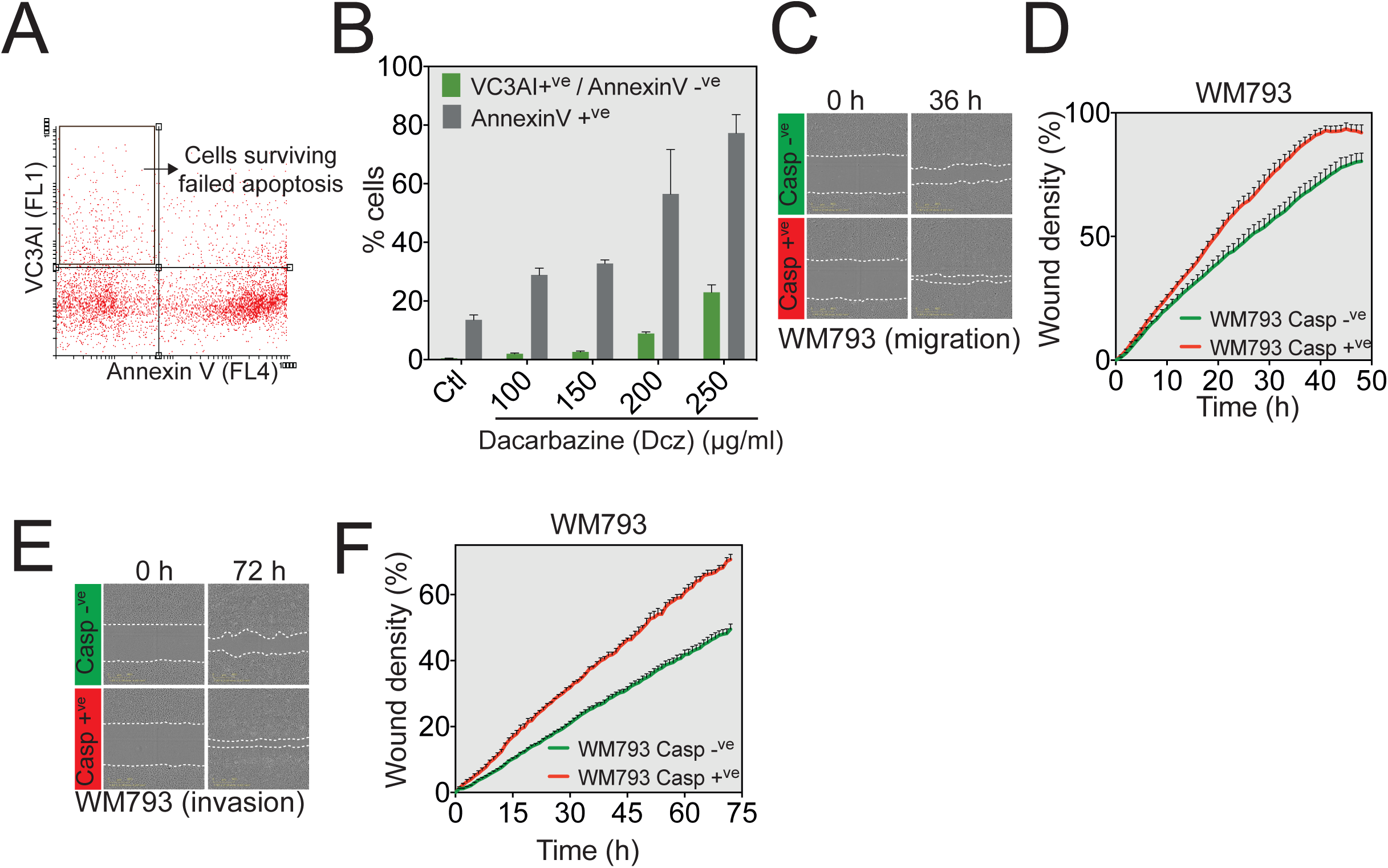
Related to Figure 5. **A**. Gating strategy used to sort and further characterize WM793 cells undergoing dacarbazine (Dcz)-induced failed apoptosis (VC3AI^+ve^/AnnexinV^−ve^) in WM793 cells. **B**. WM793 cells were first treated with different concentrations of dacarbazine for 48 hours. Cells were then stained with AnnexinV-Alexa647 and analyzed by flow cytometry according to their VC3AI (FL1) expression. **C**. Monolayer of WM793 cells treated with Dcz and sorted for VC3AI^+ve^/AnnexinV^−ve^ markers were wounded and photographs were taken immediately after wound induction and 36 hours later (migration assay). **D**. Measurement of the migratory capacity of WM793 cells through the analysis of wound area recovery using the IncuCyte imaging software. **E**. Representative images of the wound in a layer of WM793 having displayed caspase activation or not after 200 µg/mL dacarbazine treatment (invasion assay). **F**. Quantification of the invasive capacity of WM793 cells through the analysis of wound area recovery using IncuCyte imaging software.

**Supplementary Figure 6.**
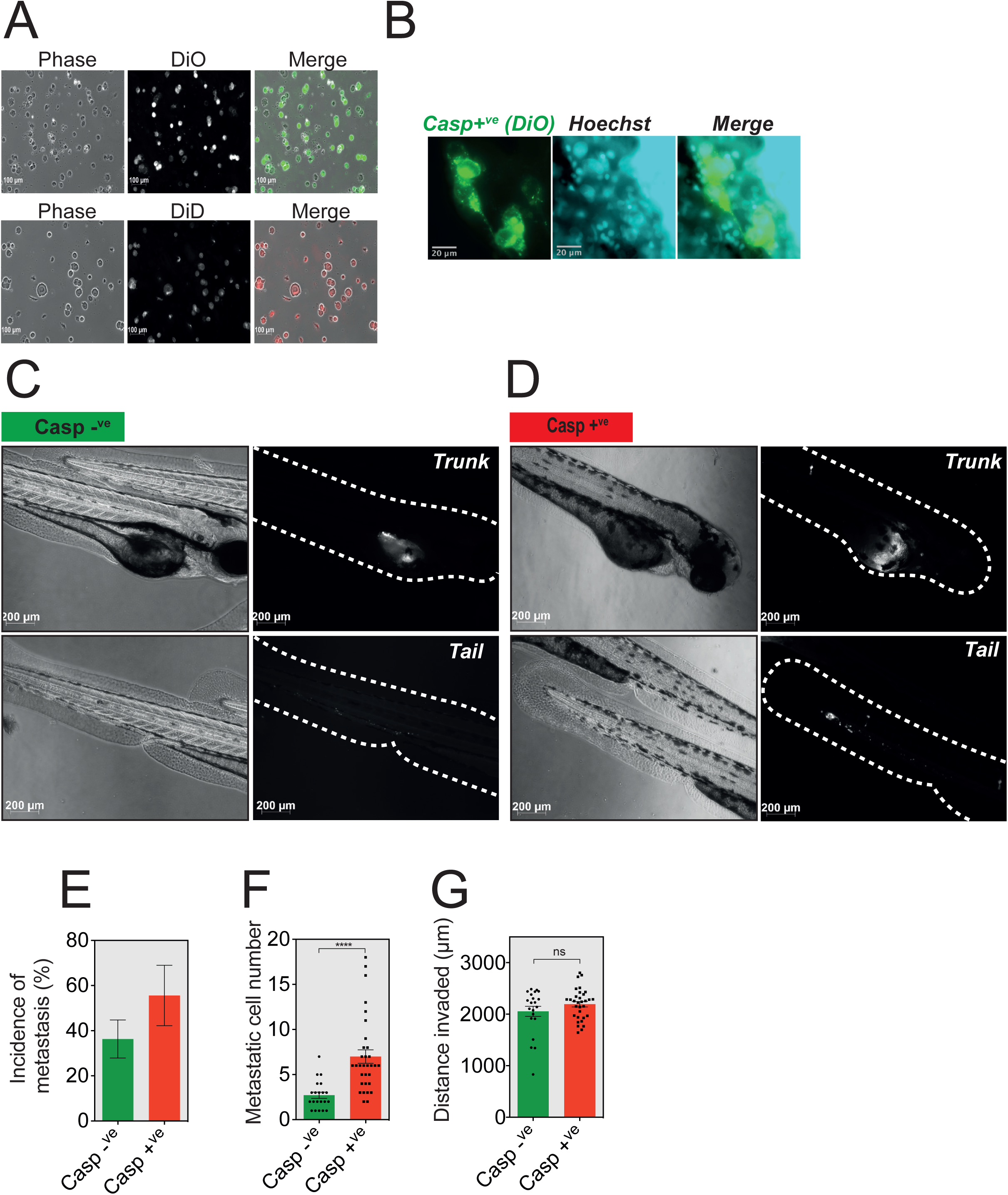
Related to Figure 6. **A**. Validation of DiO and DiD staining in WM852 cells. **B**. Invaded WM852 in the caudal blood vessels are stained with vital Hoechst and imaged for nuclear integrity. **C**. DiO-labeled Casp^−ve^ (**C**) and Casp^+ve^ (**D**) WM852 cells were injected as described in Figure 6 A and representative epifluorescence images are shown for both perivitelline homing and caudal blood vessel invasion of cancer cells. **E-G**. Quantification of metastasis incidence (**E**), metastatic cell number per embryo (**F**) and the distance invaded from the vitellus by either Casp^−ve^ or Casp^+ve^ WM852 cells (**G**).

**Supplementary Figure 7.**
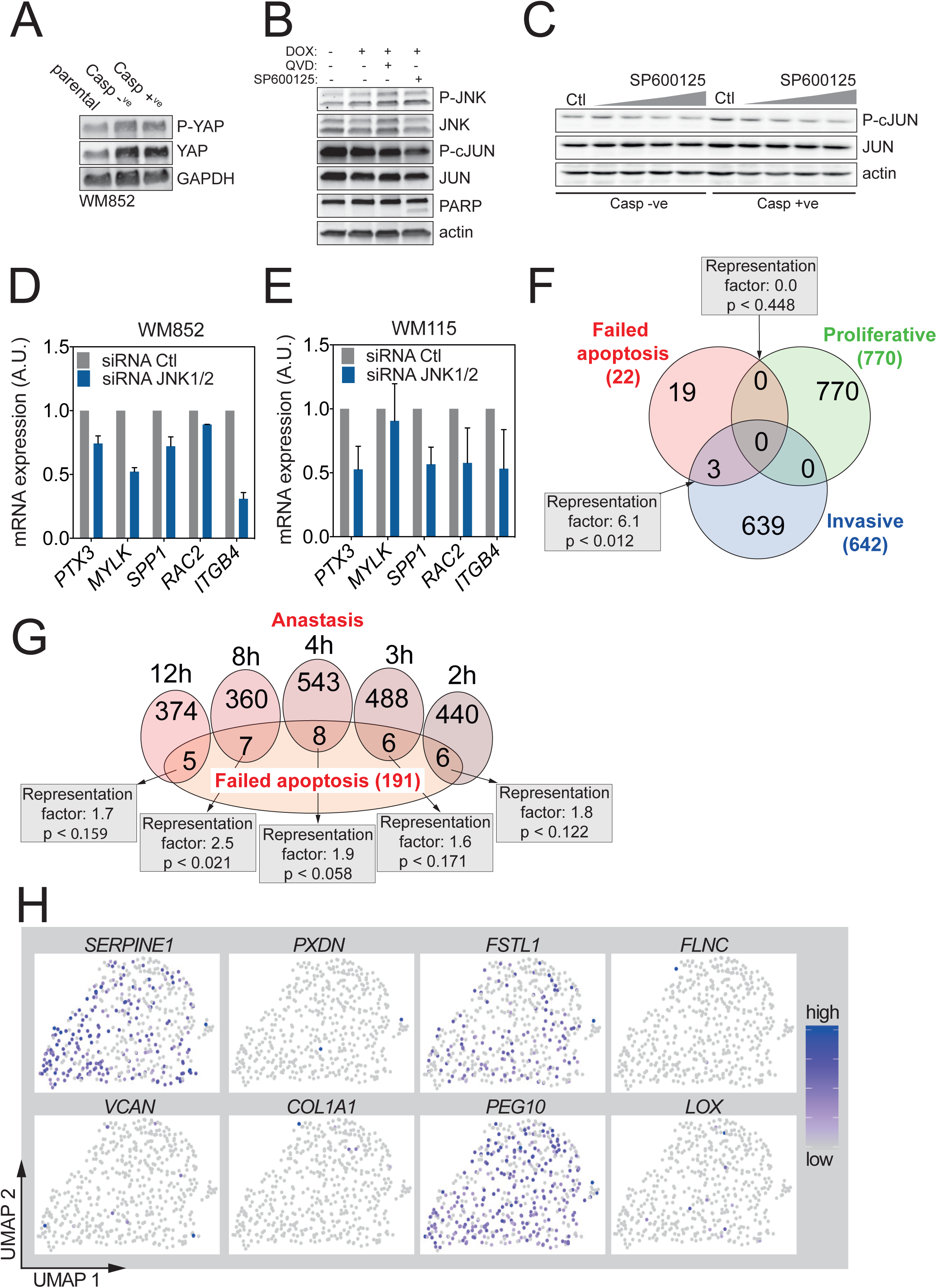
Related to Figure 7. **A**. Western blot analysis of phospho-YAP in parental, Casp^−ve^ or Casp^+ve^ WM852 cells. Actin was used as loading control. **B**. Western blot analysis of phospho-JNK and phospho-cJUN in WM852 cells treated once with 250 ng/mL of doxycycline in the presence or absence of QVD (20 μM). Actin was used as loading control. **C**. Dose-dependent effect of SP600125 (concentrations range: 20, 40, 60 and 80 μM) of JUN phosphorylation. **D-E**. Gene expression analysis by qRT-PCR in parental WM852 (**D**) and WM115 (**E**) cells following JNK1/2 knockdown. **F**. Venn diagram detailing overlaps between the 22-genes failed apoptosis signature in WM115, the proliferative and invasive Verfaillie melanoma signature. **G**. Venn diagram highlighting the overlaps between the failed apoptosis gene signature in WM852 cells and the anastasis signature at different time points. **H**. Projection plot of the failed apoptosis score for the top 8 genes of the failed apoptosis signature in each of the 464 malignant cells of a metastatic melanoma tumor.

